# Strain variation identifies a neural substrate for behavioral evolution in *Drosophila*

**DOI:** 10.1101/2025.08.15.670615

**Authors:** Anna Ryba, Philipp Brand, Rory T. Coleman, Yarden Greenfeld, Yael N. Tsitohay, Florian Hollunder, Katharine Keller, Murtaza Hathiyari, Tianyi Wei, Paolo Emilio Barbano, Vanessa Ruta

## Abstract

Sexual selection acts on heritable differences within species, driving the parallel diversification of signal production in one sex and behavioral responses in the other. This coevolution implies that sensory preferences are themselves variable traits, yet the neural basis of such variation remains unclear. Here, we identify striking strain-specific differences in *Drosophila melanogaster* male mate preferences that arise from differential sensitivity to heterospecific female pheromones. We map this variation to an ascending inhibitory pathway targeting a central circuit node known to dynamically pattern courtship. Inhibitory circuits thus emerge as a key locus for shaping mate discrimination via transient suppression of a male’s pursuit. Our findings highlight how variation within sensory circuitry serves as a substrate for selection, fueling the evolution of reproductive barriers between species.

## Introduction

Across the animal kingdom, reliance on sex has driven the emergence of fantastic anatomic and behavioral diversity, selecting for traits that reinforce reproductive isolation. Successful mating often depends on select channels of communication between conspecifics, requiring the coordination of signal production by one sex with behavioral responses by the other. Although mate recognition is thought to evolve through selection on variable traits within a species, the neural mechanisms—specifically, how heritable differences in brain circuitry give rise to divergent mating behaviors—remain largely unexplored.

*Drosophila* courtship offers a powerful window into how mate recognition systems evolve. While females are believed to decide with whom to mate, males decide whom to court^1^. In the wild, these mating decisions unfold on rotting fruits where individuals from many species congregate—ephemeral and interactive social landscapes in which males must distinguish conspecific females from morphologically similar flies^2,3^. In these chaotic sensory environments, female pheromones serve as essential cues for mate recognition^4–7^. Across the *Drosophila* genus, the long-chain cuticular hydrocarbons carried by females have rapidly and repeatedly diversified^8^, allowing males to use them as chemical signatures of sex and species identity. Comparative analyses of closely related species have revealed that as female pheromones diversify, male behavioral responses have evolved in tandem mediated by changes in both the peripheral detection of pheromones and their integration by central circuit nodes controlling a male’s sexual arousal^8–13^. Yet how these pheromone processing pathways vary within a species remains unclear.

Here, we use *D. melanogaster* strains^14,15^ —genetically and geographically distinct populations within the species—to shed light on the neural substrate of sexual selection by identifying heritable variation in the circuits that mediate mate choice. We demonstrate that males from different strains display striking variation in the fidelity of mate discrimination, due to differential behavioral sensitivity to heterospecific pheromones. Comparison of a highly promiscuous and highly selective strain reveals that differences in pheromone sensitivity map to changes in an ascending inhibitory pathway that impinges on P1 neurons, a central circuit node known to dynamically modulate the gain of visuomotor circuits controlling courtship pursuit. Inhibition of P1 neurons by heterospecific pheromones is thus able to render males transiently ‘blind’ to the presence of an inappropriate mate and trigger them to redirect their courtship pursuit towards other more suitable targets. Notably, components of the same pheromone pathways have been identified as sites of change between closely related *Drosophila* species^10,11^, suggesting how heritable strain variation may be linked to the evolution of species-specific mate preferences. Together, our work suggests a circuit logic for mate discrimination in *Drosophila*, where alterations in pheromonal inhibition can drive the evolution of mate choice without compromising a male’s sexual arousal or courtship drive, reinforcing the existence of modular circuits as a substrate in the rapid diversification of mating systems.

## Results

### Male selectivity varies across wild-derived strains

Both *D. melanogaster* and its sister species, *D. simulans* independently migrated from their ancestral range in south-central Africa to become globally distributed^16–19^. Whereas in most members of the *D. melanogaster* clade, males selectively pursue their conspecific females^20^, *D. melanogaster* males have been reported to court related but reproductively incompatible species, such as *D. simulans* females^21–23^. To explore whether male mate discrimination is a variable trait in *D. melanogaster,* we tested the courtship preferences of a panel of 11 inbred lines collected from sites in sub-Saharan Africa, as well as two European lines, an isofemale line from the Neotropics (Panama) and two common laboratory strains, Canton-S (Ohio, USA) and Oregon-R (Oregon, USA) (**Figure 1A**). Preference was measured by offering a single male access to both a *D. melanogaster* and a *D. simulans* female and quantifying courtship towards each using a combination of relative orientation and distance. Across our panel, male courtship preferences varied in a graded manner, revealing variation in the fidelity of mate discrimination (**Figure 1B**). Males from some strains were highly discriminating and upon encountering their conspecific, faithfully followed and sang to her for the majority of the assay despite occasional interactions with the *D. simulans* female (**Figure 1C**, left; **Movie S1**). Males from other strains, in contrast, were far more variable in their courtship preferences, with some individuals displaying sustained pursuit of either the *D. melanogaster* or *D. simulans* female while others alternated between them (**Figure 1C**, right; **Movie S2**). Males from promiscuous strains displayed highly variable preference indices when assayed repeatedly across days (**Figure S1A**), suggesting that an individual male arbitrarily divides his time between the conspecific and heterospecific female during a given assay, giving rise to a broad distribution of preference indices. Promiscuity in mate choice therefore does not appear to reflect the idiosyncratic preferences of individual males but instead represents a strain-specific phenotype. Notably, preference among some African strains derived from a single collection site (e.g. Gikongoro, Rwanda) varied as much as among those isolated from opposite sides of the continent, aligned with evidence that African *D. melanogaster* maintains high genetic diversity with little population structure over vast geographic territories^16^. In comparison, strains collected outside of Africa were generally promiscuous, suggesting that globally distributed *D. melanogaster* share this behavioral trait.

**Figure 1.**
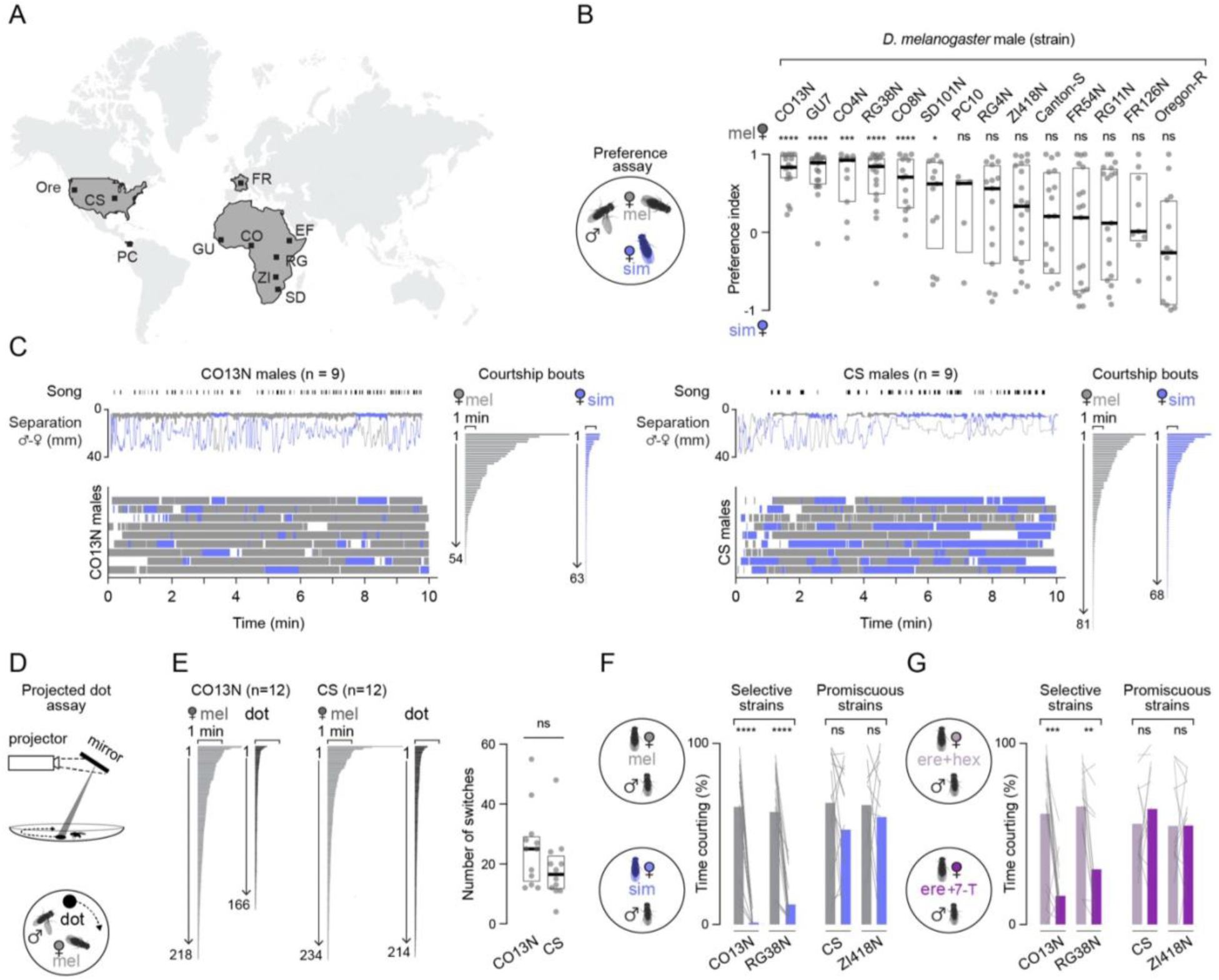
Male preference varies across *D. melanogaster* strains based on their differential responses to 7-T. (**A**) Collection locations for strains tested in the preference assay. (**B**) Courtship preference index (PI, see Methods) of males paired with a *D. melanogaster* and *D. simulans* female with individual preferences (grey dots), median (bold line), and inter-quartile range (IQR, box) shown for each strain. Six out of 14 strains tested showed a preference (one sample t-test, null hypothesis of mean PI = 0). (**C**) Courtship of selective CO13N (left) and promiscuous CS (right) males in the preference assay. Top, representative assays with bouts of song and separation between the male and the *D. melanogaster* (grey) or *D. simulans* (blue) female plotted over time. Bouts of courtship to each female indicated by a thicker line overlaid on the separation plot. Bottom, bouts of courtship by male to each female plotted over time as a raster. Courtship bouts for all assays (n=9) ordered by duration and numbered. (**D**) Schematic of the projected dot assay (see Methods). (**E**) Comparison of CO13N and CS male behavior in the dot assay. Left, bouts of courtship from all assays toward the *D. melanogaster* female (grey) and the projected visual stimulus (dot, black) ordered by duration and numbered. Right, number of times CO13N and CS courting males switched between targets in a single assay (grey dots), median (bold line), and 95% confidence interval (CI, box). (**F-G)** Percent time spent courting for selective (CO13N, RG38N) or promiscuous (CS, ZI418N) males paired with a single *D. melanogaster* (grey) and *D. simulans* (blue) female (F) and a single control *D. erecta* female treated with hexane, the carrier solvent (light purple) or one perfumed with 7-T (dark purple) (G, see Methods) with means (colored bars) and individual males (lines) shown. ns, p > 0.05; *, p < 0.05; **, p < 0.01; ***, p < 0.001; ****, p < 0.0001. Details of statistical analyses and sample sizes are given in Table S1.

Given the propensity of promiscuous males to switch between courtship targets (**Figure 1C**, right), we considered whether they have a greater tendency to become visually distracted by the motion of another female passing their field of view. However, males from both highly selective (CO13N) and highly promiscuous (CS) strains displayed a broad range of preference indices when offered two conspecific females (**Figure S1B**), suggesting that susceptibility to visual distraction is a shared trait. To assess whether a purely visual distractor could interrupt male pursuit, we examined the courtship dynamics of males offered a choice between a conspecific female and a high-contrast 3 mm dot projected onto the floor of the assay chamber that rotated at a constant angular velocity (**Figure 1D**). Both selective CO13N and promiscuous CS males were indifferent to this two-dimensional stimulus in isolation and rarely courted it (**Figure S1C**). However, once aroused by the presence of a *D. melanogaster* female—presumably by sampling her pheromones—males of both strains altered their behavior toward the visual stimulus and began to pursue and sing to it (**Movies S3-4**). While CO13N and CS males both preferentially courted their conspecific over the dot, their tendency to chase it nonetheless frequently caused them to abort their pursuit of the female, replicating the erratic bouts of courtship they displayed as they alternated between two conspecific females (**Figures 1E**, **S1D-E**). In fact, selective CO13N males showed a lower preference for the female over the dot than their promiscuous CS counterparts, due to the fact that they switched to pursue the dot more frequently (**Figures S1D-E**). Therefore, while the presence of a conspecific female promotes arousal and alters how both selective and promiscuous males engage with their visual environment, strain-specific variation in courtship preference cannot be attributed to differences in their susceptibility to visual distraction.

A key distinction between selective CO13N and promiscuous CS males was that while they engaged in a comparable number of close-range encounters with *D. simulans* females, selective males rapidly terminated their pursuit, displaying only brief bouts of courtship (**Figure S1F**). We therefore reasoned that sensory or behavioral feedback signals –such as the distinct pheromones carried by *D. simulans* females— may curtail continued pursuit by selective males. While *D. melanogaster* females produce 7,11- heptacosadiene (7,11-HD), a sex-specific diene compound that serves as a potent promoter of conspecific courtship^24^, *D. simulans* females produce 7-tricosene (7-T), the same monoene hydrocarbon carried by *D. melanogaster* males and implicated in courtship suppression^9,20,25^. To compare strain-specific responses to these chemically distinct females, we paired individuals from different promiscuous (CS, ZI418N) and selective (CO13N, RG38N) strains sequentially with a single *D. melanogaster* and *D. simulans* female and quantified courtship **(Figure 1F**). Across strains, males showed vigorous courtship towards conspecific females, consistent with 7,11-HD being a potent arousing cue. By contrast promiscuous and selective strains varied in their willingness to court a *D. simulans* female (**Figures 1F**, **S1G**): while promiscuous strains courted with the same vigor as conspecifics, selective strains rarely initiated pursuit. A similar pattern was observed when comparing the preferences of selective CO13N and promiscuous CS males offered a choice between a conspecific and a *D. yakuba* female that also carries 7-T^8^. However, both strains indiscriminately courted *D. erecta* females that lack 7-T and instead produce distinct diene hydrocarbons (**Figure S1H**). Together, these results point to differences in the behavioral response to 7-T as a potential mediator of strain-specific mate preferences.

To confirm that variation in mate preference across the broader panel of strains arises from differential sensitivity to 7-T, we perfumed target females with synthetic pheromones and examined courtship in a subset of selective and promiscuous isolates. Perfuming *D. simulans* females with 7,11-HD was sufficient to render them attractive to selective strains (**Figure S1J)**. Unexpectedly, *D. melanogaster* females perfumed with 7-T remained highly attractive to all males, (**Figure S1I**) indicating that the strong excitatory effect of 7,11-HD appears to override the inhibitory influence of 7-T when these two chemical cues are artificially encountered together. We therefore examined male responses to *D. erecta* females which lack 7,11-HD. While all males vigorously courted *D. erecta* females, perfuming them with 7-T suppressed courtship in selective (CO13N, RG38N) but not promiscuous (CS, ZI418N) strains (**Figure 1G**). *D. melanogaster* strains thus differ in their ability to use 7-T as an inhibitory cue to suppress heterospecific courtship, a behavioral difference that implicates the 7-T response pathway as a key site of variation in male mate preference.

### 7-T detection appears conserved between selective and promiscuous hybrids

Chemosensory receptor genes exhibit pronounced signatures of selection across *D. melanogaster*^26^, potentially facilitating this species’ rapid expansion into diverse ecological niches. Variation in 7-T sensitivity across strains may therefore reflect differences in peripheral pheromone detection. To identify neural correlates mediating differences in mate discrimination, we exploited the fact that F1 hybrids between the selective CO13N strain and CS transgenic lines were highly selective and maintained a strong aversion to courting *D. simulans* females (**Figure S2A**). Likewise, the analogous cross with the promiscuous CS strain bore promiscuous F1 offspring (**Figure S2A**), allowing us to introduce neurogenetic tools into selective CO13N and promiscuous CS hybrids to compare the circuit logic for heterospecific mate recognition and the underlying neural locus of variation.

In the foreleg, the neurons that detect cuticular hydrocarbons are marked by expression of the sodium channel subunit, Ppk23, an essential component of pheromone transduction machinery^10,27–29^. Selective CO13N hybrids carrying a null mutation for this receptor lost their aversion to *D. simulans* females and pursued them with the same vigor as promiscuous strains (**Figure S2B**), highlighting the critical role of Ppk23 in mediating heterospecific avoidance. Ppk23-neurons are heterogeneous but can be differentiated based on both gene expression and their functional tuning to pheromones^30^. One subset of Ppk23^+^ sensory neurons is narrowly tuned to 7,11-HD and co-expresses the vesicular glutamate transporter (VGlut) among other genes, while a second subset, implicated in detecting 7-T and other inhibitory pheromones to suppress courtship pursuit lacks VGlut expression^27,30^. We used intersectional genetics to investigate the pheromone responses of this Ppk23^+^/VGlut^-^ sensory neuron subset (**Figure S2C**) by recording the aggregate activity of their axon terminals within the ventral nerve cord as a female abdomen was brought into contact with a male’s foreleg, allowing him to sample her pheromones. Ppk23^+^/VGlut^-^ neurons responded strongly to both *D. melanogaster* and *D. simulans* females (**Figure 2A**), indicating that this subset may still represent a heterogenous population, sensitive to multiple classes of pheromones that shape courtship in an opposing manner. Nevertheless, pheromone responses appeared indistinguishable between promiscuous CS and selective CO13N hybrids, demonstrating that 7-T is robustly detected in both strain backgrounds.

**Figure 2.**
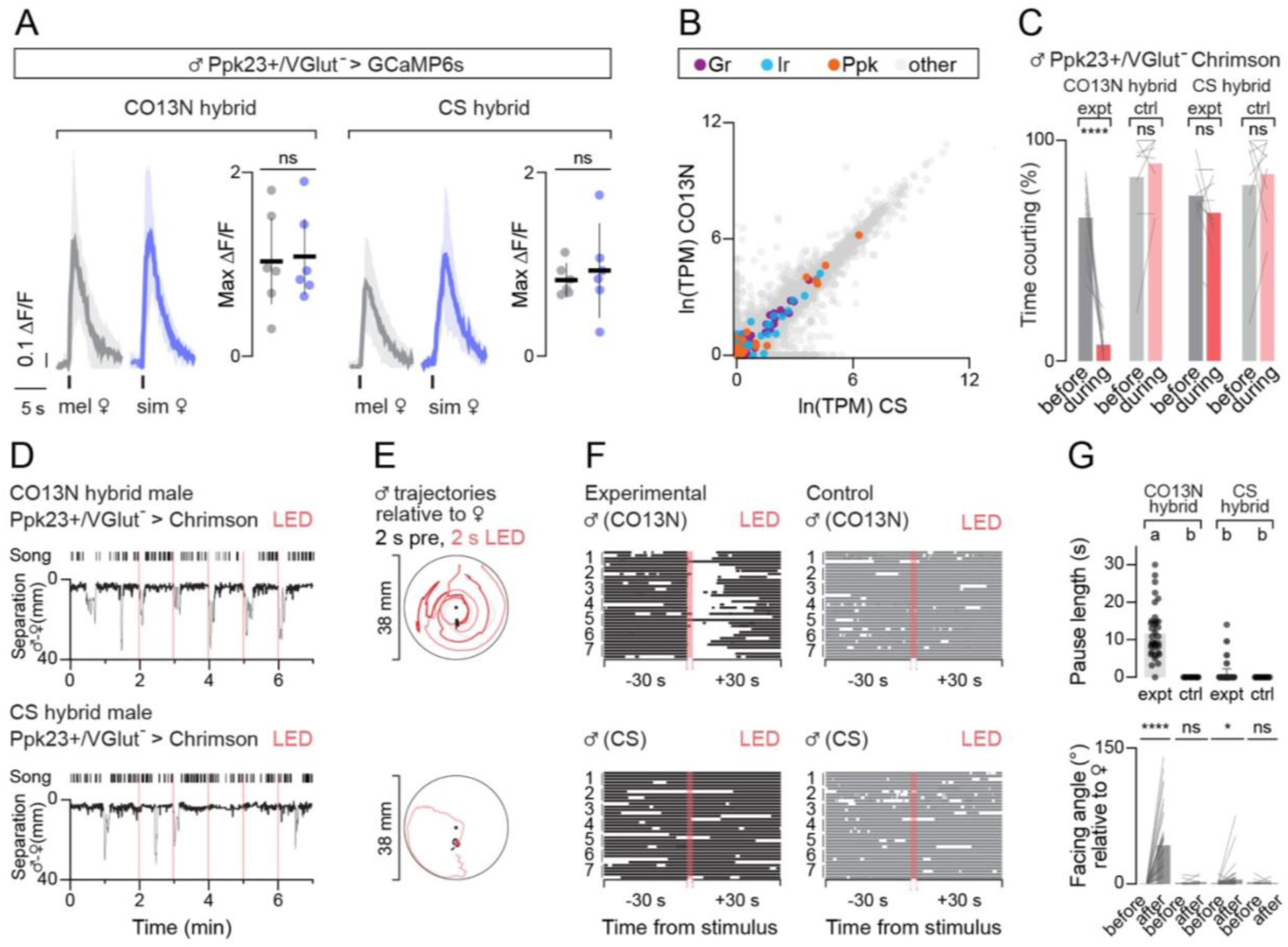
7-T detection appears conserved between selective and promiscuous hybrids. (**A**) Responses of Ppk23^+^/VGlut^-^ neurons in CO13N (left) and CS (right) hybrids evoked by tapping a *D. melanogaster* (grey) or *D. simulans* (blue) female abdomen. Left panel shows mean (Δ*F*/*F_0_*) aligned to tap (black tick) and right panel shows maximum responses. For aligned responses, shading is 95% CI. For peak responses, individuals (dots), mean (line), and 95% CI (error bars) are shown. (**B**) Relative abundance (transcripts per million, TPM) on a natural log scale of all transcripts in CS (x-axis) and CO13N (y-axis) male forelegs with gustatory receptors (GRs, purple), ionotropic receptors (IRs, blue), and pickpocket channels (Ppks, orange) highlighted. (**C-G**) Courtship toward a *D. melanogaster* female in CO13N and CS hybrids expressing Ppk23-Gal4, VGlut-Gal80 > UAS-CsChrimson raised on food with retinal (experimental animals) or without retinal (controls). (**C**) Percent time spent courting 30 seconds prior to (grey) and during 30 second optogenetic stimulation (red) with means (colored bars) and individual males (lines) shown. (**D**) Representative CO13N (top) and CS (bottom) hybrid males subjected to five 2-second pulses of optogenetic stimulation. Bouts of wing song and separation plotted over time with bouts of courtship indicated by a thicker line overlaid on the separation plot and timing of stimulus marked by red bars. (**E**) Trajectories of males in (D) relative to the female’s position in the 2 seconds prior to (black) and 2 seconds during optogenetic stimulus (red) for each of the five stimuli. (**F**) Courtship bouts 30 seconds before, during, and 30 seconds after the five optogenetic stimuli in (D) for all males. The five stimuli for each male are shown and aligned to time of stimulation for experimental (left, black) and control (right, gray) CO13N hybrid (top) and CS hybrid (bottom) males. (**G**) Top, duration of pause in courtship evoked by activation of Ppk23-Gal4, VGlut-Gal80 > UAS-CsChrimson for all males in (F) with individual stimulation epochs (dots), mean (bars), and 95% CI (error bars) shown. Bottom, male facing angle relative to the female (degrees) in the 30-second window prior to and after all stimuli in (F) with individual stimulation epochs (lines) and mean (bars) shown. ns, p > 0.05; * < 0.05, ****, p < 0.0001, groups that are not significantly different from one another (p > 0.05) after multiple comparisons indicated by shared letter codes. Details of statistical analyses and sample sizes are given in Table S1.

The molecular logic for 7-T detection remains to be fully defined, with several classes of chemoreceptors implicated in pheromone recognition and signal transduction^12,25,27,31,32^. To examine potential differences in chemosensory gene expression across strains in an unbiased manner, we performed RNA sequencing of the forelegs from two promiscuous strains (CS, RG11N), a selective strain (CO13N), and a selective hybrid (CS/CO13N). We isolated RNA from just the distal tarsal segments of the foreleg (**Figure S2D**), given that this region is densely packed with the Ppk23^+^ sensory neurons essential to mate recognition^33^ (**Figure S2B**) and its surgical removal is sufficient to render selective males promiscuous (**Figure S2E**). None of the four main classes of *Drosophila* chemosensory genes (gustatory receptors (GRs), odorant receptors (ORs), ionotropic glutamate receptors (IRs), and pick-pocket channels (Ppks) displayed significant differences in abundance between selective and promiscuous strains (**Figure 2B**). A small number of genes were significantly up- or down-regulated (82 of 13,635 genes with > 1 log-fold change in relative abundance, adjusted p-value < 0.05) between promiscuous and selective strains (**Table S1**), but none were associated with chemosensory detection or neuronal signaling. Moreover, pairwise nucleotide diversity across all expressed chemosensory gene transcripts (GRs, IRs, Ppks, ORs, OBPs, and CSPs) was low, independent of whether comparisons were made across selective (CO13N) and promiscuous (CS or RG11N) strains or two promiscuous strains (CS and RG11N) (**Figure S2F**).

Thus, transcript nucleotide sequence and transcript expression data, along with functional evidence (**Figure 2A**), suggest that behavioral differences between selective and promiscuous strains are unlikely to reflect differences in 7-T detection but rather represent variation in the downstream circuits that integrate and interpret this chemical cue. Concordant with this, optogenetic activation of the Ppk23^+^/VGlut^-^ population—which bypasses pheromone detection—recapitulated the distinct behavioral sensitivity of selective and promiscuous hybrids to 7-T. Sustained activation of CsChrimson-expressing Ppk23^+^/VGlut^-^ neurons led to a pronounced suppression of male courtship towards a conspecific female in CO13N but not CS hybrids (**Figures 2C, S2G**). Transient stimulation (2 seconds) of Ppk23^+^/VGlut^-^ sensory neurons—intended to mimic the discrete sampling of pheromonal cues that males perform during natural courtship—further highlighted strain-specific differences. Brief activation of this sensory population disrupted the close pursuit of CO13N hybrid males, triggering them to reorient and circle the chamber before re-encountering the female and resuming courtship (**Figures 2D-G**; **Movie S5**). By contrast, CS hybrid males were largely indifferent to Ppk23^+^/VGlut^-^ sensory neuron activation and courted through the optogenetic pulses (**Movie S6**). Ppk23^+^/VGlut^-^ neurons thus display similar sensitivity to 7-T across strains, but their activation differentially interrupts courtship in selective but not promiscuous males.

### Differences in P1 neuron tuning to 7-T mirror strain selectivity

Given that strain variation does not appear to originate at the sensory periphery, we searched for differences within the central brain circuits that process pheromonal cues. P1 neurons form an integrative node in a male’s courtship circuitry that guide his ongoing performance of the courtship ritual by dynamically modulating the visuomotor circuits orchestrating pursuit^34–38^. In *D. melanogaster* males, P1 neurons are excited by *D. melanogaster* female pheromones, aligned with evidence that they trigger an enduring state of sexual arousal to promote pursuit of conspecifics^30,39^. Across *Drosophila* species, while P1 neurons play a conserved role in driving courtship, they display divergent pheromone tuning^10,11^, underscoring significant evolutionary plasticity in the detection and processing of pheromones across species and suggesting a substrate for how male mate preferences may diversify in parallel with changing female pheromone profiles.

To assess whether the chemosensory tuning of P1 neurons differs among *D. melanogaster* strains, we recorded P1 responses in tethered males as they walked on an air supported ball and were presented with conspecific and heterospecific females to tap and taste, replicating the discrete sensory sampling that occurs during courtship encounters^10,11^. P1 neurons were robustly activated by *D. melanogaster* females in both selective CO13N and promiscuous CS hybrids (**Figure 3A**), consistent with the vigorous courtship males of parental strains displayed towards conspecific females (**Figures 1F, S1G**). By contrast, the pheromones of a *D. simulans* female evoked negligible responses in the P1 neurons in either strain (**Figure 3A**), despite the strong attraction that promiscuous males display to these heterospecific females. These results support behavioral evidence that neither the presence of 7-T nor activation of the Ppk23^+^/VGlut^-^ sensory neurons that detect this pheromone promotes courtship in promiscuous strains (**Figures 2C, S1I**). Rather, 7-T appears to shape mate choice by transiently suppressing the courtship pursuit of selective strains, presumably by countering other excitatory signals, such as vision, that promote and perpetuate a male’s arousal (**Figures 1D, 2D-G**). Indeed, excitatory pheromones like 7,11- HD are dispensable in *D. melanogaster* in the light when males have access to vision^9^. Therefore, to assess whether 7-T can attenuate excitatory inputs to the P1 neurons, we presented males with perfumed *D. erecta* females, allowing us to directly relate the sufficiency of 7-T to suppress courtship at the behavioral level to the suppression of P1 neuron activity. While mock-perfumed *D. erecta* females elicited comparable responses in the P1 neurons of both selective and promiscuous hybrids (**Figures 3B, 1G**), perfuming them with 7-T strongly suppressed tap-evoked excitation only in selective CO13N hybrids (**Figure 3B**), providing a direct neural correlate for the divergent behavioral responses to this pheromone observed between strains.

**Figure 3.**
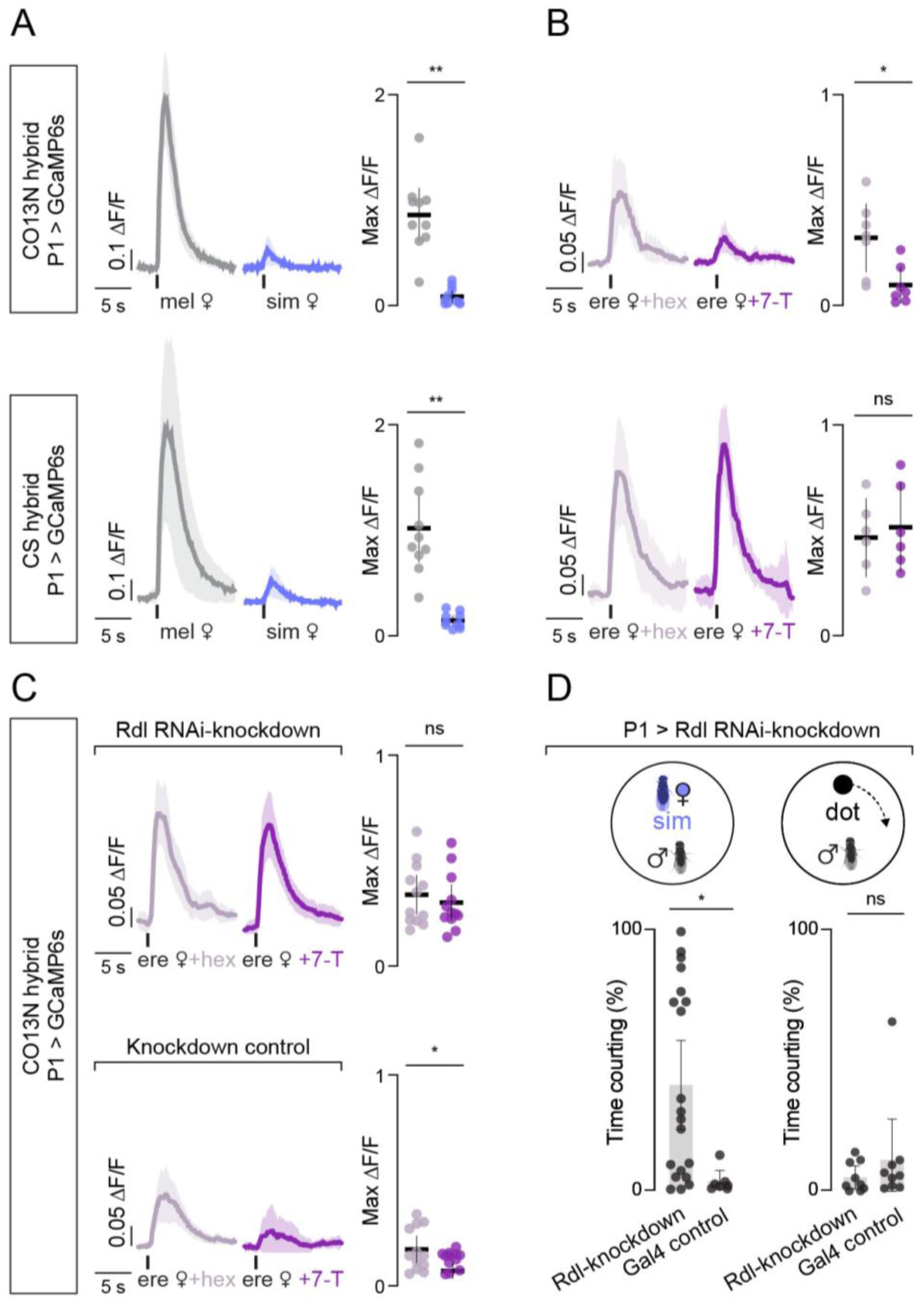
Differences in P1 neuron tuning to 7-T mirror strain selectivity. (**A**) P1 neuron responses in CO13N (top) and CS (bottom) hybrids evoked by tapping the abdomen of a *D. melanogaster* (grey) or *D. simulans* (blue) female. Left panels: mean Δ*F*/*F_0_* traces (thick line) aligned to tap (black tick) with 95% CI shading. Right panels: peak responses showing individual data points (dots), mean (line), and 95% CI (error bars). (**B**) P1 responses as in (A) but evoked by tapping the abdomen of a control *D. erecta* female treated with hexane (light purple) or one perfumed with 7-T (dark purple). (**C**) As in (B), but in CO13N hybrids expressing UAS-Rdl.RNAi in P1 neurons (top) or Gal4 controls (bottom). (**D**) Time spent courting a *D. simulans* female (left) or a projected dot (right) in CO13N hybrids in which Rdl has been knocked down or in Gal4 controls. Dots indicate individuals; bars show mean; error bars show 95% CI. ns, p > 0.05; *, p < 0.05; **, p < 0.01; ****, p < 0.0001. Details of statistical analyses and sample sizes are given in Table S1.

Courtship-promoting P1 neurons thus appear differentially sensitive to 7-T in selective and promiscuous strains, suggesting that quantitative variation in inhibitory drive to this central node may underlie differences in mate preference. To explore this possibility, we therefore sought to weaken inhibitory signaling in the P1 neurons of selective hybrids using RNA-interference to knockdown Rdl, a GABA_A_ receptor subunit highly expressed in this population^30,40^. We found that this manipulation abolished the sensitivity of P1 neurons to 7-T-mediated inhibition in CO13N hybrids, resulting in equivalent functional responses to both 7-T- and mock-perfumed *D. erecta* females (**Figure 3C**). Attenuating inhibition in the P1 neurons of CO13N hybrids is thus sufficient to phenocopy the pheromonal tuning of promiscuous males. Consistent with this neural phenotype, Rdl knockdown rendered selective CO13N hybrid males significantly more promiscuous and willing to court *D. simulans* females (**Figure 3D**, left). Nevertheless, knockdown animals remained largely indifferent to a projected visual target rotating on the floor of the assay chamber (**Figure 3D**, right), underscoring that reduced inhibition at the level of P1 neurons does not inherently promote a male’s arousal but rather specifically alters how chemical cues shape his courtship pursuit.

### Differences in an inhibitory chemosensory pathway determine strain selectivity

Together, our data point to the pathways that link 7-T sensing neurons to P1 neuron activity as a potential site of diversification driving variation in *D. melanogaster* mate selectivity. Ppk23-neurons form part of an ascending circuit that bifurcates to convey balanced excitation and inhibition onto P1 neurons and tune their responses to pheromones^30,39^. Given the importance of inhibitory inputs to mate discrimination, we focused on the ∼30 mAL GABAergic neurons^41^ which comprise one branch of this pheromone circuit (**Figures 4A**). mAL neurons are thought to counterbalance the excitatory input to P1 neurons elicited by the taste of a conspecific female^39^, and indeed, functional imaging of their axon terminals within the lateral protocerebral complex (**Figure 4A, S3A**) revealed strong responses to a *D. melanogaster* female target that were comparable between hybrid strain backgrounds (**Figure 4B**). By contrast, responses to a *D. simulans* female were far weaker in promiscuous CS hybrids than selective CO13N hybrids (**Figure 4B**). In both hybrid strains, pheromone responses were abolished in ΔPpk23 mutants (**Figure S3B**), consistent with the essential role that Ppk23^+^ sensory neurons play in mediating 7-T detection.

**Figure 4.**
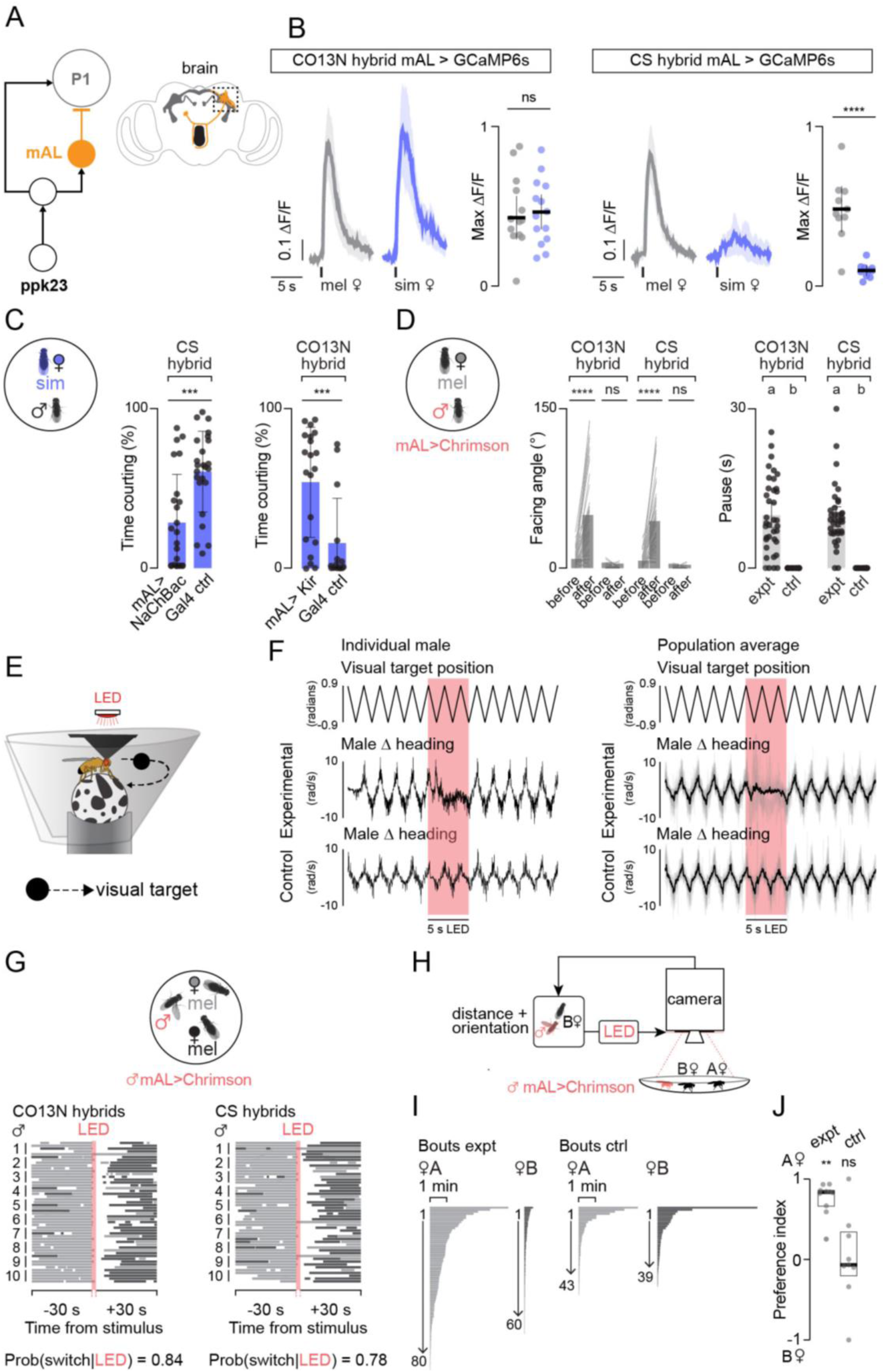
Differences in an inhibitory chemosensory pathway determine strain selectivity. (**A**) Diagram (left) and anatomic cartoon (right) summarizing proposed chemosensory circuit^30,39^ connecting Ppk23-neurons in the foreleg to P1 neurons (grey) in the brain via inhibitory mAL neurons (orange). (**B**) mAL neuron responses in CO13N (left) and CS (right) hybrids evoked by tapping the abdomen of a *D. melanogaster* (grey) or *D. simulans* (blue) female. Left panels: mean ΔF/F_0_ traces aligned to tap onset (black tick) with 95% CI shading. Right panels: peak responses showing individual data points (dots), mean (line), and 95% CI (error bars). (**C**) Time spent courting a *D. simulans* female in mAL > UAS-NaChBac and Gal4 control CS hybrid males (left) or in mAL > UAS-Kir and Gal4 control CO13N hybrid males (right). Individuals (dots), mean (bars), and 95% CI (error bars) shown. (**D**) Courtship of a *D. melanogaster* female in mAL > UAS-CsChrimson CO13N and CS hybrids, raised on food with retinal (experimental animals) or without retinal (controls). Left: Male facing angle relative to the female (degrees) during 30 second before and after optogenetic stimulation; individual trials (lines) and mean (bars) shown. Right: duration of pause in courtship evoked by 2 second optogenetic stimulus with individual trials (dots), mean (bars), and 95% CI (error bars) shown. (**E**) Schematic of tethered courtship paradigm used for mAL activation. (**F**) Left: position of the dot in front of the tethered male (top), Δ heading angle (degrees/second) for a representative tethered mAL > UAS-CsChrimson male, raised on food with retinal (middle), and a representative control male, raised on food without retinal (bottom) with optogenetic stimulus period marked in red. Right: Same as left but for all trials, with individual traces shown as grey and mean in black. (**G**) Courtship bouts by mAL > UAS-CsChrimson CO13N (left) and CS (right) hybrid males toward one of two *D. melanogaster* females 30 seconds before, during, and after a 2 second optogenetic stimulus. For each stimulus presentation, the bout immediately prior to stimulation is in light grey and subsequent bouts are colored by female identity to highlight switches in courtship after stimulation. (**H**) Schematic of the closed loop optogenetic system. (**I**) Courtship bouts ordered by duration and numbered for mAL >UAS-CsChrimson hybrid males raised on food with retinal (left) or controls lacking retinal (right). (**J**) Preference index for experimental and control males from (I) courting two *D. melanogaster* females in the closed-loop assay with individuals (grey dots), median (bold line), and inter-quartile range (IQR, box) shown. ns, p > 0.05; **, p < 0.01; ***, p < 0.001; ****, p < 0.0001, groups that are not significantly different from one another (p > 0.05) after multiple comparisons indicated by shared letter codes. Details of statistical analyses and sample sizes are given in Table S1.

Equivalent responses at the level of peripheral Ppk23^+^ neurons are thus transformed into differential mAL responses, indicating that mAL-mediated inhibition may shape mate discrimination by controlling inhibitory input to the P1 neurons. Indeed, constitutively depolarizing mAL neurons via expression of the bacterial sodium channel, NaChBac^42^, rendered promiscuous males more selective (**Figure 4C**, left), while constitutively hyperpolarizing this population via expression of the inward rectifying potassium channel, Kir2.1^43^, made selective males more promiscuous (**Figure 4C**, right). Bidirectionally manipulating the excitability of mAL neurons thus predictably alters male behavior, pointing to changes in this inhibitory node as a potent regulator of mate discrimination.

Concordant with this idea, brief (2 second) optogenetic activation of mAL neurons (**Figure S3C**) temporarily interrupted male courtship in both hybrid backgrounds, triggering males to abort their pursuit and reorient from the female, before rapidly resuming courtship upon reencountering her several seconds later (**Figures 4D**, **S3D**). Notably, while activating Ppk23+ peripheral sensory neurons triggered divergent behavioral responses in selective and promiscuous hybrids (**Figures 2D-G**), optogenetic activation of mAL neurons uniformly interrupted male courtship regardless of strain background. Strain differences in 7-T sensitivity thus appear to be localized to variation in mAL neuron responses or the ascending pathways that link Ppk23^+^/VGlut^-^ sensory neurons to this inhibitory node, such that bypassing their pheromonal input results in comparable control of courtship across strains.

### mAL-mediated inhibition interrupts courtship pursuit

Our data suggest that mate choice emerges through an ongoing evaluative process as males repeatedly sample the pheromones of a potential partner and based on this discrete sensory assessment either sustain or abort their pursuit. How does pheromone-mediated inhibition influence this decision? P1 neurons dynamically modulate the gain of the visuomotor circuits that control courtship pursuit, such that a male is indifferent to small moving targets like a projected dot (**Figure S1C**)–or even a female–until he becomes sexually aroused, whereafter the fidelity and vigor of his pursuit is correlated with P1 activity on a moment-to-moment basis^35^. mAL-mediated inhibition of P1 neurons is therefore expected to acutely decrease the gain of visual responses. Indeed, in contrast to the two-dimensional projected dot (**Figure 1D**), tethered males perceive a dot sweeping back and forth on a visual panorama in the center of their field of view as a female and will faithfully track and sing to it (**Figure 4E**). Transient optogenetic activation of mAL neurons triggered males to abort their pursuit of the visual target, despite its continued presence in their visual field (**Figure 4F**). Males resumed their pursuit of the target immediately after cessation of the optogenetic stimulus, underscoring that mAL activation transiently renders males behaviorally insensitive or ‘blind’ to the visual target.

Given that *Drosophila* courtship occurs in dense groups in the wild^2^, we reasoned that the momentary interruptions in courtship evoked by mAL activation could increase a male’s likelihood of switching his visual pursuit to a different female, thereby allowing him to more broadly sample prospective sexual partners. To explore this idea, we transiently activated mAL neurons in males paired with two conspecific females, which reliably interrupted courtship in both CO13N and CS hybrids, triggering them to temporarily abort their pursuit before reinitiating courtship, typically of the first female they encountered (**Movies S7-8)**. Given that mAL activation drove males to reorient, this led them to frequently switch the target of their pursuit to the other female (Probability of switching ∼0.80) (**Figure 4G**). If the original female was encountered first, however, males nevertheless resumed vigorous courtship indicating that they do not appear to retain a memory of which target they were courting at the onset of activation. Courtship vigor remained high throughout the assay, underscoring that mAL activation transiently disrupts a male’s pursuit without dampening his courtship drive (**Figure S3E**).

mAL mediated inhibition is thus poised to dynamically shape courtship pursuit, triggering males to disengage from heterospecific females while remaining persistently aroused and visually sensitive to the presence of potential conspecific mates in the area. To demonstrate that transient bouts of mAL-mediated inhibition could generate the strict mate preference observed in selective strains, we offered a promiscuous male two identically reared conspecific females in a closed-loop preference assay (**Figure 4H**). Prior to the start of the assay, one female was arbitrarily designated as a fictive ‘*D. simulans’* and each time the male was positioned close enough to sample her pheromones, a brief optogenetic mAL stimulus was delivered. Males rapidly aborted their pursuit of the fictive heterospecific and switched to pursuing the other female that triggered no inhibitory feedback (**Figures 4I, S3F-G**; **Movie S9**). As a result, males preferentially courted control females (**Figure 4J**) mimicking the pursuit dynamics of naturally selective strains (**Figures 1B-C**). Mate preference, then, does not arise from an inherent attraction to the control female, but from her failure to trigger inhibition, underscoring how transient pheromonal cues are sufficient to steer a male’s courtship toward appropriate mates.

## Discussion

Female pheromones are known to be critical determinants of species-specific mate recognition in *Drosophila*^4–7^, requiring that as these sexual signals diversify, male behavioral responses must coevolve. Here we leveraged natural strain variation in *D. melanogaster* as a window into the evolution of mate recognition systems. We uncovered widespread variation in mate selectivity across *D. melanogaster* strains that maps onto a shared behavioral phenotype: sensitivity to the pheromone 7-T. Comparative analyses of the underlying neural circuitry revealed that this variation arises from differences in the strength of ascending inhibitory input to the P1 neurons that dynamically control courtship pursuit. These findings identify inhibitory pheromone-processing circuits as a key locus for generating intraspecific behavioral variation that could serve as a substrate to ultimately fuel the evolution of mate preferences. Indeed, the differences we observed across strains bear striking parallels to differences in the pheromone circuits between *D. melanogaster* and its sister species^10,11^, where alterations in the balance of excitation and inhibition onto the P1 neurons enable the same pheromone to promote courtship in one species and suppress inappropriate pursuit in another. Examining differences in mate preference across *D. melanogaster* strains thus provides insight into the diversity of circuit variants available to natural selection—revealing the substrate from which species-specific behaviors originate.

Why is promiscuity so broadly maintained in *D. melanogaster*? *Drosophila* opportunistically colonize rotting fruits—ephemeral food sources where many individuals of different species congregate, creating a complex social backdrop for courtship and copulation. Widespread promiscuity could be retained in *D. melanogaster* due to local adaptation (e.g. in response to the relative abundance of receptive conspecific and heterospecific females). Alternatively, it could be maintained as a trade-off with other reproductive or survival needs or tied to *D. melanogaster’s* history as a human commensal. Notably, all the cosmopolitan isolates we tested were promiscuous (**Figure 1B**), raising the possibility that this trait arose or became accentuated by the expansion of *D. melanogaster*—alongside the spread of agriculture —from eastern sub-Saharan Africa into Eurasia around the end of the last Ice Age^16,17^. Genetic data suggest that this single event introduced a bottleneck to globally distributed cosmopolitan forms of the species^44^ that may have biased them toward promiscuity, either through a relaxation in selection for strong preference or through neutral processes like drift. Cosmopolitan *D. melanogaster* retain relatively low genomic diversity and display several phenotypic differences compared to African strains^14,45,46^, aligned with the idea that expansion and/or adaptation to new environments strongly shaped the prevalence of certain traits in the species.

In *Drosophila*, an important factor for a male’s reproductive success is the need to compete for access to receptive conspecific females, which are thought to be rare on the crowded food patches where courtship unfolds^47^. Males may therefore benefit from sampling among multiple potential mates rather than investing too heavily in the pursuit of any single individual. In general, aroused males appear to “try their luck” and indiscriminately pursue moving visual targets, as underscored by the fact that they alternate between pursuit of 2-D inanimate objects and conspecific females (**Figures 1E, S1D-E**). In selective strains, contact pheromones contribute to correcting a male’s inherently error prone visual pursuit, triggering him to abort courtship of heterospecific females and redirect his efforts towards other visual targets. Notably, however, a male’s courtship vigor remains sustained even as he repeatedly samples inhibitory pheromones from these inappropriate mates. Thus, the circuits controlling sexual arousal appear segregated from those mediating mate discrimination, suggesting that pheromone responses are free to diversify without impairing courtship vigor. By leveraging naturally occurring strain variation, we therefore shed light onto how the rapid diversification of male mate preferences may emerge from selection on modular chemosensory circuits.

Like many behavioral traits shown to vary within a population, the genetics of mate preference appear complex and likely multigenic^48^. Hybrid crosses among strains in our panel showed variable levels of courtship toward *D. simulans* females that could not be predicted from their parental phenotypes, supporting the idea that promiscuity may likewise have a complex genetic basis (**Figure S3H**).

Nevertheless, we found that promiscuity maps to a simple behavioral trait shared across different strains —decreased sensitivity to 7-T (**Figure 1G**)—pointing to ascending inhibitory chemosensory pathways as a repeated site of diversification. Consistent with this, we could render selective males more promiscuous by altering pheromone processing at multiple levels within the 7-T circuit: preventing peripheral 7-T detection (**Figure S2E**), decreasing mAL excitability (**Figure 4E**), or reducing P1 sensitivity to inhibitory inputs (**Figure 3D**). Different perturbations at multiple distinct nodes in the same chemosensory circuit thus lead to a convergent behavioral phenotype, reinforcing the idea that the widespread promiscuity we observed across different strains may arise from distributed neural and genetic loci. More generally, our results suggest how the multigenetic architecture of behavioral traits may nevertheless belie a relatively simple neural logic for behavioral diversification in which small differences at various levels of a sensory circuit orchestrate coherent changes in behavior.

**Table 1.**
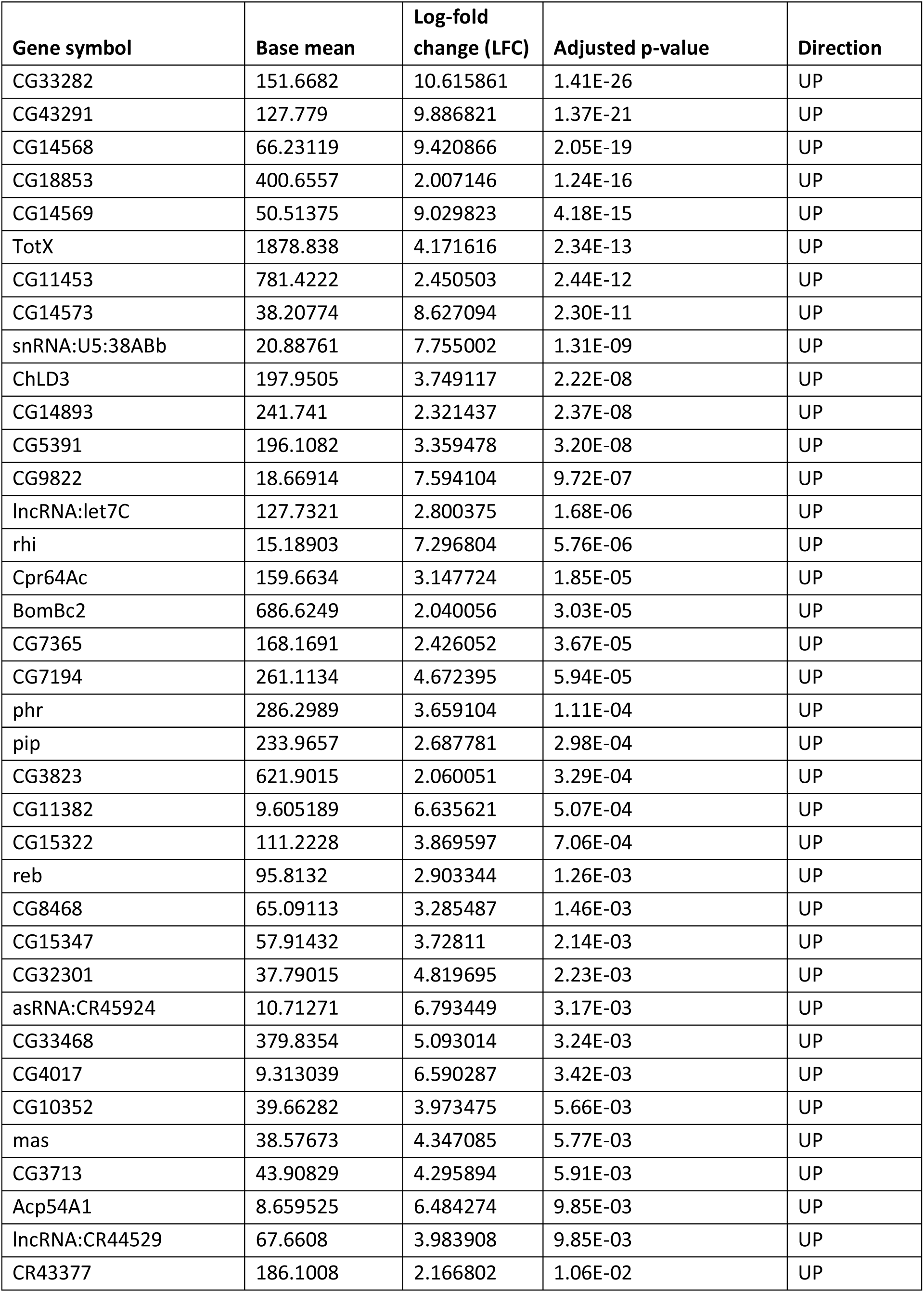
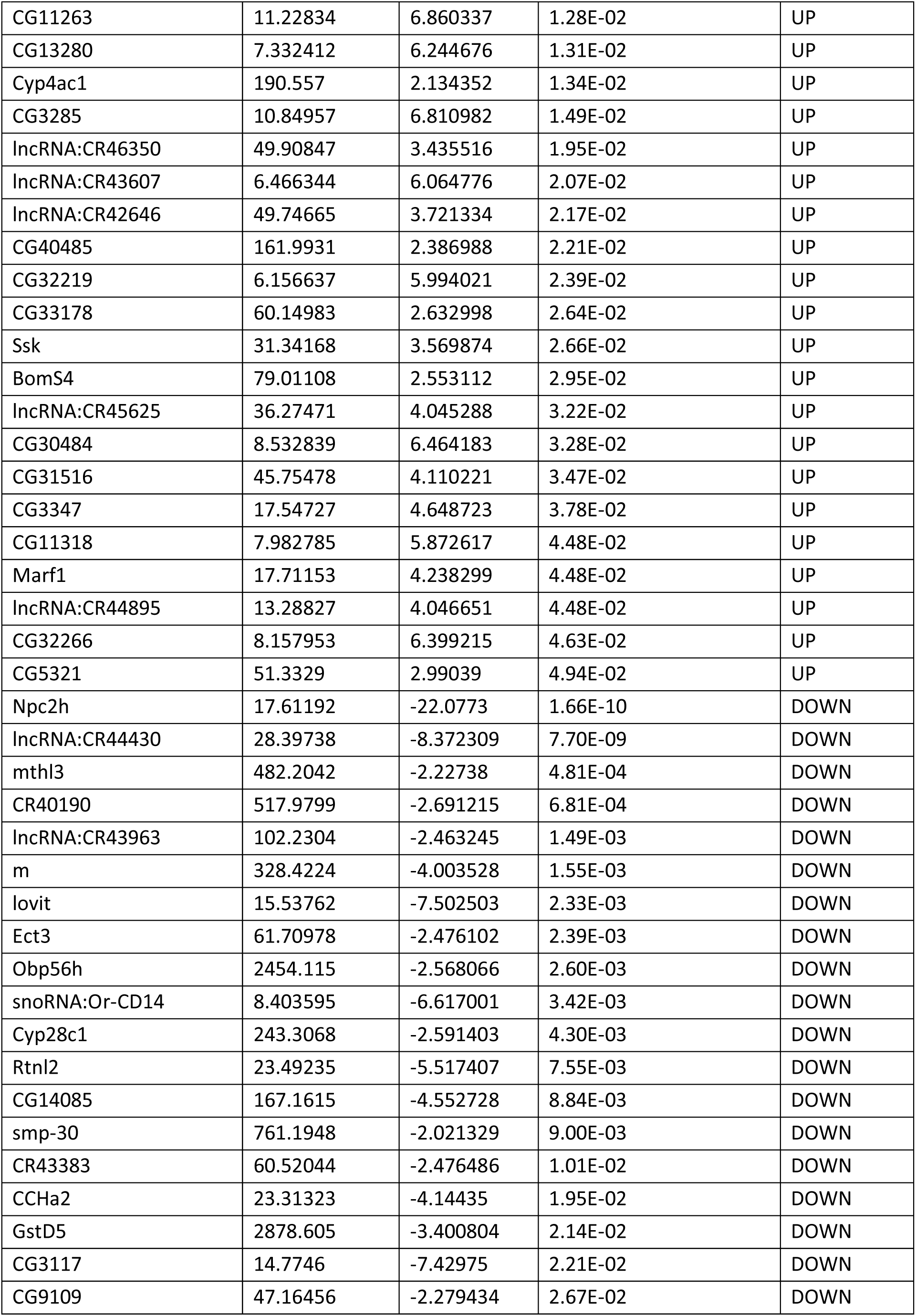
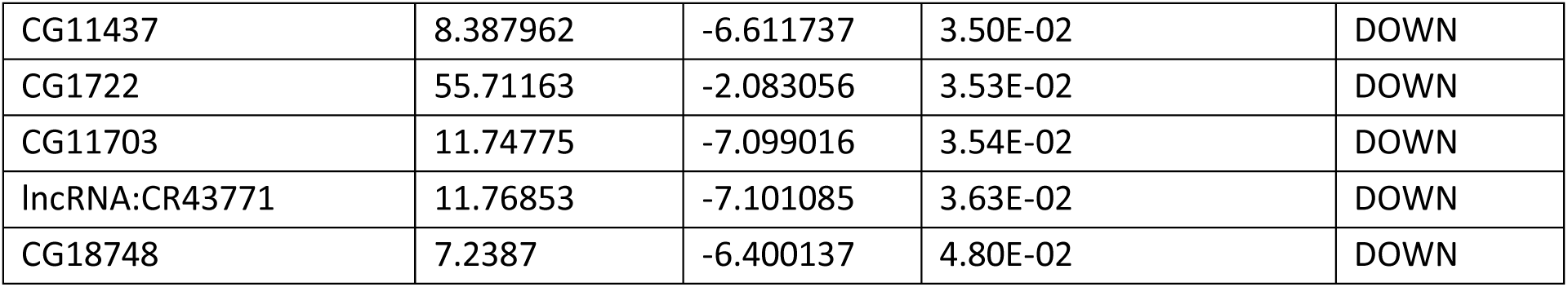
Differential gene expression in forelegs of selective vs. promiscuous males. Lists all genes that are significantly (p < 0.05) down- or upregulated (-1 < LFC < 1) between foreleg samples from selective (CO13N) and promiscuous (CS and RG11N) males according to differential expression analysis performed in DeSeq2. Additional information about level of expression, magnitude of difference in expression, significance, and direction of change in expression also defined.

## Acknowledgments

We thank C. S. McBride, B. Noro, B. Datta, M. Shahandeh, L. Vosshall, D. Kronauer, J. Rhee and all members of the Ruta laboratory for valuable discussion and comments on the manuscript; J. Pool for providing African and European fly stocks, information regarding their origin, and discussion about genomic diversity; L. Zhao for providing the PC10 line; R. Li for providing code for fly behavior quantification; M. Galperin and P. Stock for previous work on optogenetics behavioral setups; the Rockefeller University Precision Instrumentation Technologies facility for access to fabrication equipment and assistance with PCB design and fabrication; the Rockefeller University Genomics Core for library prep, RNA-sequencing, and quality control; the Rockefeller University Bioinformatics Resource Center for assistance with alignment and quantification of RNA-sequencing reads; the Rockefeller University Bioimaging Resource Center for access to a confocal microscope and assistance with collecting foreleg images. This work was supported by a Kavli Neural Systems Institute Fellowship and NIH NIGMS grant K99GM151471 (PB); a Helen Hay Whitney Foundation Fellowship and NIH NIGMS grant K99GM141319 (RTC); an NSF GRFP (YG); a Boehringer Ingelheim Fonds Ph.D. Fellowship (FH); an NIH NINDS grant R35NS111611 (VR) and the Simons Foundation Collaboration for the Global Brain (VR). VR is an Investigator of the Howard Hughes Medical Institute.

## Author contributions

AR designed and performed experiments, analyzed data, and interpreted the results, with input from VR. PB and FH designed the projector system for free behavior. FH and KK performed projector experiments. PB performed and analyzed open loop tethered behavior experiments. YG collected RNA-seq data and assisted with analysis. RTC and YNT assisted with initial preference assays and analysis. TW, with input from PEB, developed the code for real-time pose estimation and tracking for closed loop optogenetic experiments. MH designed hardware and wrote code to integrate with real-time tracking. AR and VR wrote the manuscript with input from all authors.

## Competing interests

Authors declare that they have no competing interests.

## Resource availability

Lead contact: Further information and requests for resources and reagents should be directed to and will be fulfilled by the lead contact, Vanessa Ruta

## Experimental models and subject details

Flies were housed at 25°C and 50-65% relative humidity on a 12h light:12h dark cycle. *D. melanogaster* Canton-S (stock number 64349), Oregon-R (25211), UAS-GCaMP6s (42746, 42749), R71G01-Gal4 (39599), R25E04-Gal4 (49125), UAS-CsChrimson.mVenus (55134), UAS-mKir2.1.tdTomato (600376), UAS-NaChBac.EGFP (9466), UAS-VGlut-Gal80 (58448), UAS-Rdl.RNAi(8-10) (89903), R25E04-p65.AD (68874) were obtained from the Bloomington *Drosophila* Stock Center. Inbred isofemale lines RG38N, RG11N, RG4N, CO13N, CO4N, CO8N, GU7, SD101N, SD105N, ZI251N, ZI418N, FR54N, FR126N, EF8N, EF43N, EF101N, SP39N, SP107N (John Pool, University of Wisconsin-Madison); isofemale line PC10 (Li Zhao, Rockefeller); Fru-Gal4.DBD (Barry Dickson, Janelia); Ppk23-Gal4 (Kristin Scott, UC Berekley); ΔPpk23 mutants were generated in a previous study^10^. *D. yakuba* Ivory Coast (14021-0261.00) and *D. erecta* (14021-0224.01) were obtained from the National *Drosophila* Species Stock Center (formerly Cornell Stock Center). *D. simulans* (Ruta lab, origin unknown). See **Table S2** for detailed genotypes by figure. All experimental males were isolated from females, housed in new food vials with other males, and used 2- 4 days post-eclosion; all test females were virgin, housed with other females, and used 2-4 days post-eclosion, unless otherwise noted.

## Methods details

### Back-crossing drivers

To generate selective and promiscuous animals for neurogenetic experiments, necessary transgenes were first crossed into a single animal. If two or more transgenes were located on the same chromosome, they were recombined using standard *Drosophila* genetics techniques, then male parents carrying all transgenes were crossed to virgin CO13N or CS females. Pilot experiments showed that whether the transgenes came from a male or female parent did not affect behavior, but this parental scheme a) eliminated w^-^, a widespread marker on the X-chromosome and b) minimized the probability of transgene loss from recombination, which functionally does not happen in *Drosophila* males. When hybrids were generated using new sets of transgenes, courtship was assessed for a set of 3-6 males in single pairings with *D. simulans* females to ensure that the experiment was comparing behaviorally selective and promiscuous strains.

### Courtship assays and analysis

All assays were conducted at 25°C, 46% relative humidity and 0 to 3 hours after lights on, except for optogenetics experiments in which males had been reared in the dark and were used throughout the day. Males and virgin females were added by direct aspiration from the food vial. Unless otherwise stated, assays were performed in 38mm diameter, 3mm high circular, sloped-walled chambers, backlit using a white light pad (Logan Electric), and recorded from above using a Point Grey FLIR Grasshopper USB3 camera (GS3-U3-23S6M-C: 2.3 MP, 162 FPS, Sony IMX174, Monochrome) fitted with a 2.8-10mm CS mount lens (FLIR, LENS-28C2-V100CS).

#### Preference assays

Video recording was started before the addition of flies to the courtship chamber (and subsequently trimmed from raw videos before tracking). A male and two female flies (A and B, order of introduction was noted) were aspirated into the chamber, at which point the 10-minute assay commenced. Preference index = (time the male spent courting female A - time spent courting female B) / total time spent courting during this 10-minute period. (Thus, preference is a fraction between -1 and 1 where 0 means the male split his time evenly between both females.) *Criteria for exclusion –* Males that courted for less than 20% of preference assays were not included, since at that point a comparison between extremely brief bouts of courtship with one female or another became somewhat meaningless. Discarded assays made up <10% of all trials.

#### Single pairing assays

A single virgin female and male were aspirated into the courtship chamber, video recording was started, and a 10-minute assay was recorded. Courtship index (expressed as a percentage) = total time courting / total time in the assay. Where paired experiments are indicated in the results, the following procedure was followed: males were paired first with one female, the video recording was ended, the female was removed via aspiration, the next female was added via aspiration, and the next video was started. Pilot experiments showed that the order of pairing did not affect courtship index. However, when experiments included a conspecific female, males were paired with her second to minimize the number of trials that needed to be discarded due to copulation during the assay. For the tarsi ablation assays, males were ice-anesthetized 24 hours prior to the start of the experiment and had the distal four tarsal segments on both forelegs removed with a scalpel. Males were then returned to a food vial to recover in isolation before testing. *Criteria for exclusion –* If males copulated less than 10% into the assay, that assay (or that set of paired assays) was not included. Discarded assays made up <5% of all trials. Otherwise, if the male copulated >10% into the assay, the total time in the assay was taken from the assay start to the moment of copulation and all calculations were performed as above.

#### Free behavior visual stimulus projection assays

Our projector system design was inspired by previous setups using stationary ^49^ and moving visual cues^50^ in a behavioral arena. We performed our assays in the 38mm slope-walled assay chambers described above, coated with white self-adhesive film to enhance reflectance, and maximize contrast of the projected stimulus. Using a Lightcrafter 3010 micro projector (Texas Instruments), we projected a circular dot (∼2.5mm diameter) moving at a constant angular velocity circling the arena in a 30mm diameter once every 3 seconds directly onto the assay chamber surface. Tester males were aspirated into the assay chamber after which the visual stimulus projection started, and recording commenced for 10 minutes using a Basler acA3088-57uc camera with an 8mm/F1.8 lens (Edmund Optics). If pairs of flies were used, the female was aspirated first after which the visual stimulus and recording started directly followed by aspiration of the male into the chamber.

#### Perfuming

Protocol was adapted from previous perfuming methods^9^. Virgin females were placed in a 1mL amber vial with 7,11-HD (7(Z), 11(Z)-heptacosadiene, 10 mg/mL Cayman Chemicals #100462-58-6) or 7-T (7(Z)-tricosene, 10 mg/mL Cayman Chemicals # 9000313) dissolved in carrier solvent (pure hexane) that was then evaporated off under air. Control, hexane vials were prepared by evaporating off pure hexane under air. Females were placed in a vial for perfuming or control treatment, respectively, vortexed gently, and left to recover for 30 minutes, then transferred to a fresh food vial to groom for 1-4 hours before testing.

#### Note on RNAi

UAS-Rdl.RNAi(8-10) was made by Ron Davis’s lab (Scripps, Florida) and obtained from Bloomington. This construct does not have a scrambled control, so Gal4 controls were used.

#### Free behavior optogenetics

All optogenetic animals were reared on standard sugar-yeast (SY) food in vials covered in aluminum foil. Upon eclosion, experimental animals were transferred to foil covered vials for 2-4 days on food containing 400μM all-trans-retinal (Sigma R2500-10MG); in contrast, control animals were transferred to fresh foil covered vials containing standard sugar-yeast (SY) food. For video recordings of behavioral experiments, background illumination was diffused by placing the courtship chamber on top of a white acrylic sheet (220mm by 220mm by 3mm) suspended 6cm over a custom LED panel. The panel consisted of 9 white (6500K Cool White, A007-CG2H765S4, LEDdynamics) and 9 near-IR (850nm IR LED, LZ1-10R602-0000, ams-Osram AG) starboard LEDs in evenly spaced 3-by-3 matrices that provided a source of light for the flies to court and for clear video, respectively. LEDs were mounted over a layer of thermal paste on a custom aluminum panel (152.4mm by 114.3mm by 8mm) bolted on a 24cm-by-24cm Thorlabs breadboard to improve heat dissipation. Each LED channel was connected in series and driven by a custom dual channel LED driver developed by Nicholas Belenko at the Gruss Lipper Precision Instrumentation Technologies (PIT) using Pulse Width Modulation (PWM). This driver connected to an Arduino Uno to allow finer control over light intensity and was externally powered by a 30V-5A benchtop power supply (Naweisz, NP6005). The white LEDs were driven at 30% maximum intensity, which in pilot experiments was sufficient to elicit courtship indices comparable to those in full light but which didn’t activate CsChrimson from either optogenetic driver (Ppk23-Gal4/VGlut-Gal80 or 25E04-p65.AD/Fru-Gal4.DBD) used here. The IR LEDs were set to strongly illuminate the video image but at relatively low power, which prevented any burnouts. Behavior was recorded using a Grasshopper camera fitted with near-IR long pass filter (Machine Vision Direct, LP800-49) to avoid overexposure of the video image during LED-on periods.

For Ppk23^+^/VGlut^-^ optogenetic activation, experimental and control animals were Ppk23-Gal4, VGlut- Gal80 > UAS-CsChrimson-tdTomato hybrids. For mAL optogenetic activation, animals were 25E04-p65.AD, Fru-Gal4.DBD > UAS-CsChrimson-tdTomato hybrids, as this intersection labels a sparser subset of neurons than 25E04-Gal4 and pilot experiments showed that it eliminated motor effects at a range of LED intensities. Single male flies were loaded, followed by single female flies. Pairs were left together until the experimenter observed the first bout of courtship (defined by the first observation of wing extension); video recording was then started and experiments were conducted according to the following protocols using custom software for data acquisition and instrument control. Chronic activation: two minutes baseline (dim white light), 30 seconds constant LED illumination (60% maximum intensity, 530nm Precision LED Spotlight with Uniform Illumination–PLS-0530-030-S, Mightex Systems), one minute baseline after light offset. Brief activation: two minutes baseline, then the following was repeated five times: 2s LED stimulation (60% maximum intensity, 10 Hz, 50% duty cycle), one minute baseline after light offset. In all assays, courtship index was quantified for each period separately and if index for the full assay was needed, indices were summed across chunks.

#### Head-fixed optogenetics

For head-fixed, open loop mAL activation assays, animals were 25E04-p65.AD, Fru-Gal4.DBD > UAS-CsChrimson-tdTomato CS hybrids (CO13N hybrids were not tested, as they behaved indistinguishably from CS hybrids in free behavior assays, see **Figures 4D, 4G, S4D-E**). Males were briefly anaesthetized on CO_2_ and tethered to a custom-milled head plate similar to those used in previous studies^35,51^ using UV-curable glue (Bondic). After recovery, flies were transferred onto a virtual reality setup described in Sten et al. 2021, adapted from Weissman, J. and Maimon, G. In brief, the tethered flies were positioned on an air supported 6.35mm diameter ball, whose movements were imaged, and the male trajectories were instantaneously tracked using FicTrac^52^. A circular visual stimulus (∼28° diameter, mimicking the angular size of a female fly 2mm away from the male) was projected onto a conical screen surrounding the male, moving in a symmetric 100° arc at constant angular velocity and size. Red LEDs (530nm Precision LED Spotlight with Uniform Illumination–PLS-0530-030-S, Mightex Systems) were mounted on a custom panel close to the fly’s head. Experiments were performed according to the following protocol: male behavior was recorded for 15 seconds at baseline without the visual stimulus, after which the visual stimulus was presented for the remaining assay. Once the visual stimulus turned on, behavior was recorded for 30 seconds, then red LED was activated for 5 seconds. This stimulus paradigm was repeated 3 times and the assay stopped after a final 30 second period without LED activation.

#### Real-time closed-loop optogenetics

This experiment required: custom hardware, software for real-time pose estimation and tracking, and a pipeline for integrating hardware and software components. The system was built in collaboration with the Rockefeller University Data Science Platform.

Behavior was conducted in the same 38mm chambers described above and backlit with the same panel. A ring of 8 red starboard LEDs (CREEXPE2-COL-X, LED Suppy) connected in series were affixed to a custom aluminum ring (153.58mm diameter, 10mm thick, 10mm fins) suspended around the camera ∼8cm above the surface of the behavior chamber for optogenetics. The arena was centered about the ring to ensure even optogenetic illumination across the chamber and LEDs were driven by a single channel LED driver operated in essentially the same way as the dual channel driver. Since we were observing long term preference behaviors in these experiments, we assumed the delay introduced due to serial communication with the Arduino to be negligible. External power was supplied similarly as in the backlighting setup by a 30V-5A benchtop power supply (MAISHENG, QW-MS305D). Behavior was recorded from above by an industrial camera (Basler ace acA1920-150um Monochrome USB 3.0 Camera) fitted with a near-IR long pass filter (Machine Vision Direct, LP800-49) that streamed frames directly to a software pipeline for camera-tracker-optogenetics integration.

Data acquisition and control was implemented on a single custom developed machine from Exxact Corporation (Processor: Intel i9-14900K 24 cores, GPU: NVIDIA GEFORCE RTX 4090 - 24GB GDDR6X GPU, RAM: 32GB DDR5 4800M/Ts, OS: Ubuntu 22.04). The software for running the closed-loop experiments was written in Python because of the availability of various open-source libraries and ease of interfacing with the code. Additionally, we took advantage of multi-processing in Python to reduce the overhead time for each iteration of the closed loop pipeline. The main process controlled all steps that needed to happen serially in any given iteration of the experiment: image acquisition, pose-estimation and tracking (see below), algorithmic calculations for controlling the red optogenetic LEDs, and two-way communication with the Arduino for executing optogenetics. Saving acquired images, saving raw data, and a GUI with a real-time stream of the video data with animal identity overlaid were all implemented in separate multi-processes. Additionally, software allowed for online control to change the hardware and software parameters before the start of the experiment to ensure clear image acquisition necessary for maintaining identity across the acquired frames. At the start of the experiment the software connected to the camera and began streaming video at 30 Hz, executing an iteration of the pipeline for each acquired frame. Each iteration of the pipeline was executed in ∼11ms (∼9ms for pose estimation and tracking, ∼2-3ms for the remaining overhead).

Pose tracking consisted of three steps: fly pose estimation, identity tracking, and track correction. For each frame captured by the camera, the pipeline detected the positions of flies, predicted sex labels, and estimated the positions of keypoints for each detected fly using YOLOv11-Pose^53^. The identity tracking framework was adapted from ByteTrack^54^, a multi-object-tracking algorithm associating high-confidence and low-confidence detection boxes with existing tracklets in two matching stages. However, detection using the standard box association method (using Intersection Over Union) often led to identity swaps: the detection boxes around courting pairs of flies frequently overlapped, making it difficult to maintain correct assignments. This issue was addressed by deploying the keypoint distance (KD) as a cost metric for track matching, leveraging keypoint estimations to incorporate both positional and orientation information. KD between a pair of detected flies in adjacent frames was calculated:

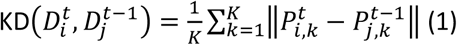

where 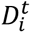 is the detected fly𝑖at frame𝑡, 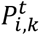 is the estimated position of keypoint 𝑘of animal 𝑖at fram 𝑡,and 𝐾 is the number of keypoints. For each tracker update, the keypoint distances were computed between detected flies in the current frame and the tracked flies from the previous frame and stored in a 2D cost matrix. The standard Hungarian algorithm^55^ was applied on the cost matrix to determine the optimal assignment with the lowest cost. Tracking consisted of two steps. First, a low distance threshold was used to associate detected flies (confidence score >0.1) with existing tracks, ensuring that all non-occluded detections were associated with their corresponding tracks, thus preserving continuous and reliable movement patterns. Second, a higher distance threshold was applied to associate remaining detections (confidence score >0.3) with unassigned tracks. This step was crucial for handling rapid fly movements: when a fly moved too fast, it was not detected in some frames. Once the fly slowed down and became clearly detectable again, its new position was sometimes far from its previous track. Re- tuning the distance threshold allowed relinking such detections, thereby ensuring that the fly’s track was not lost due to temporary occlusion.

Given that during courtship behaviors a very small number of male-female identity swaps still occurred, we implemented an *a-posteriori* identity correction mechanism that continuously tracked the sex label assigned to each track and maintained a record of the male fly’s track ID after each tracker update. If the male fly’s track ID changed for a certain number of consecutive frames, it indicated that its identity had been swapped with one of the female flies: the algorithm then reassigned the correct track IDs to the male and female flies, restoring their original identities.

Experiments were performed as follows: an experimenter loaded two *D. melanogaster* females into the courtship chamber, then added a 25E04-p65.AD/Fru-Gal4.DBD > UAS-CsChrimson CS hybrid male raised on food with retinal (experimental) or without retinal (SY food, control). Animals were left together until the male commenced courtship with either female (defined by the first observed instance of wing extension). The experiment was started within the software and run until completion. To reiterate, male position and orientation relative to each female was captured and calculated in real-time and 500ms constant red LED was delivered when he was <2mm from the arbitrarily designated female and kept her within a 60° arc in front of him. Titration experiments helped determine an intensity of 35 μW/mm^2^ for driving the optogenetic response. Video and tracking data were saved for each experiment, courtship toward each female was quantified (see Scoring courtship), and preference index was calculated (see Preference assays). In about half of the trials begun, detection using the algorithm outlined above did not happen correctly. In these instances, the experiment was terminated within the first 10 seconds by the experimenter and the software restarted. The experiment was started again, at which point detection usually succeeded and the experiment proceeded (although if detection failed again, the restart process was repeated until detection succeeded). These failures at the detection stage were because of low confidence predictions of the tracks which has been addressed in the tracking code uploaded to GitHub by reassigning identity every 120 iterations. If copulation occurred with either female before the end of the experiment, only data collected until copulation was used to calculate preference index. If copulation happened less than 20% into the 10-minute assay (before 2 minutes), the trial was discarded.

#### Scoring courtship

For all assays except those with the dot and the closed-loop optogenetics, courtship and/or wing song were scored using (x,y) position, facing angle, and wing angle data tracked using FlyTracker^56^. Courtship was defined as periods during which males were <5mm from a female and kept her within a 60° arc in front of them. Wing song (for the purposes of visualization – not a formal criterion for courtship) was defined as periods during which wings were at an angle from the main body axis. For **Figure S1F** encounters were defined by instances when males were <2mm from the *D. simulans* female and kept her within a 60° arc in front of them. For assays with the dot, occlusion of the male by the dot caused insurmountable issues for automated tracking, and courtship was instead quantified by a blinded observer and logged in BORIS^57^. For closed-loop optogenetics, online tracking data saved by the system was used to define courtship as above.

### Two-photon functional imaging

All imaging experiments were performed on an Ultima two-photon laser scanning microscope (Bruker Nanosystems) equipped with galvanometers driving a Chameleon Ultra II Ti:Sapphire laser. Emitted fluorescence was detected with either photomultiplier-tube or GaAsP photodiode (Hamamatsu) detectors. Images were acquired with an Olympus 60Å∼ 1 numerical aperture objective and collected at 512 pixel x 512 pixel resolution with a frame rate from 0.2-0.4 Hz when imaging an ROI. Saline (108mM NaCl, 5mM KCl, 2mM CaCl2, 8.2mM MgCl2, 4mM NaHCO3, 1mM NaH2PO4, 5mM trehalose, 10mM sucrose, 5mM HEPES pH7.5, osmolarity adjusted to 275mOsm) was used to bathe the brain for all imaging experiments. Fluorescence time-series were extracted using FIJI (v.2.14.0/1.54f). Ventral nerve cord and LPC preparations were performed as previously reported^10,39^. For all imaging experiments, the presentation order of stimuli was randomized and the strongest responding stimulus was presented again the end of an experiment to confirm the continued health of the experimental male.

### Imaging analysis

For peripheral imaging, an ROI was drawn in the first thoracic segment of the ventral nerve cord neuropil where Ppk23+ neurons send dense afferents. For P1 and mAL imaging, an ROI was drawn in the LPC neuropil where axonal projections from each set of neurons, respectively, were densest. For all experiments, 3-6s were recorded before the stimulus presentation to create a baseline. Twenty frames from this period were averaged to determine baseline fluorescence (F0) and ΔF/F0 was calculated as ΔF/F_0_ = (F_t_ − F_0_)/F_0_, where t denotes the current frame.

### Foreleg RNA-sequencing

Tissue preparation and RNA extraction were adapted from previous methods^58^.

#### Tissue preparation

3-8 day-old flies were CO_2_-anesthetized and kept on the pad no longer than 20 minutes. The foreleg was removed by forceps below the sex comb, transferred to DNA Lo-bind nuclease-free tubes (Fisher Scientific #13-698-790), and once 100 legs were collected (or 20 minutes had elapsed), the tube was flash-frozen in liquid nitrogen. Caution was taken during the tissue dissection and RNA extraction process to ensure that there was no contamination from other fly tissues or RNases. The CO_2_ pad, forceps, and scalpels were cleaned with 70% ethanol and RNase-away (ThermoFisher #7003) after every round of dissection. 90-100 forelegs were used for each library. Sample groups were dissected in parallel to avoid batch effects. Tubes were flash frozen in liquid nitrogen and allowed to thaw twice before dissected tissue was stored at 80°C until RNA extraction.

#### RNA extraction

RNA extraction was performed using the PicoPure Kit (ThermoFisher #KIT0204) with the following exception for homogenizing tissue: instead of lysis buffer, 300uL of TRIzol (ThermoFisher #15596018) was added to the collection tube on ice. Custom-order molecular biology grade, low-binding zirconium beads in 100mm, 200mm and 800mm were used to disrupt tissue (OPS diagnostics). An RNase free spatula (Corning #CLS3013) was used to add 1 scoop each of 100mm and 200mm beads and 100uL of 800mm beads to collection tube. Tubes were briefly spun down in a tabletop centrifuge before disruption in a TissueLyser II (QIAGEN #85300) for 2.5 minutes at 30Hz, spun down and returned to the TissueLyser II for an additional 2 minutes at 30Hz, then spun and returned to the TissueLyser II for 1 minute at 30Hz. The remaining TRIzol extraction steps were performed in a chemical fume hood according to manufacturer’s instructions: tubes stood at room temperature for 5 minutes before 48uL of chloroform:isoamyl alcohol 24:1 was added (Sigma #C0549). Tubes were hand-shaken for 30s and left to stand for 2 minutes before centrifuging at 12,000xg for 15 minutes at 4°C. The aqueous Trizol layer was then removed and added into the PicoPure column, up to 180uL at one time. Subsequent steps were performed according to PicoPure manufacturer’s instructions, including DNase treatment.

#### RNA-sequencing

Foreleg samples were run on a Bioanalyzer RNA Pico Chip (Agilent #5067-1513) to determine RNA quantity and quality then sent to The Rockefeller University Genomics Resource Center for PolyA selection and library preparation using the Nextera XT DNA library preparation kit (Illumina #FC-131-1024). Samples (3 each for CO13N, CS, and hybrid) were pooled before distributing the pool across sequencing lanes. Single-end sequencing was performed at The Rockefeller University Genomics Resource Center on a NextSeq 500 sequencer (Illumina). All reads were 1 x 75bp. Data were de-multiplexed and delivered as fastq files for each library to us and to the Rockefeller University Bioinformatics Resource Center (BRC). The BRC trimmed all reads and performed preliminary quality analysis.

#### Sequencing analysis

Analysis was performed in R. Reads from individual libraries were mapped to the BDGP *D. melanogaster* genome assembly version 6^59^ using the Subjunc aligner in the Rsubread library. For abundance visualization, raw counts were converted to TPM using Salmon^60^. Plots were generated using ggplot2 version 3.2.0^61^ in R-Studio. DESeq2 version 1.24.0^62^ was used to build differential expression models from raw counts.

To quantify sequence diversity of expressed transcripts among strains, SNPs of coding regions in the Dm6 RefSeq annotations were called based on the RNA-Seq data. The three replicates per strain were combined using samtools after which haplotypes were called in GATK’s HaplotypeCaller optimized for RNA-Seq data with the --dont-use-soft-clipped-bases flag on. Based on this data SNPs were called and combined for all strains in GATK taking only sites with a minimum coverage of 15x per strain into account. The dataset was then filtered by several metrics requiring mapping quality >40, strand bias <50, strand odds ratio <4, coverage per allele >10, and variant quality normalized by depth of coverage <1.5. All transcripts were filtered with a minimum of 95% sites called in all strains. Pairwise nucleotide diversity (π) between all strain combinations was calculated, where π=variant sites/all sites. Based on this dataset the position of all synonymous and non-synonymous SNPs between all strains for all transcripts of the chemosensory-related genes (Ppk10, Ppk15, Ppk23, Ppk25, Ppk29, Or43b, Ir25a, Ir52c, and Ir76b) in this filtered dataset were called.

### Statistical analysis

GraphPad Prism 9 was used for statistical analyses and plotting. Prior to analysis, the Shapiro-Wilk method was used to test normality, and the appropriate parametric/non-parametric analysis was used. In cases for which multiple comparisons were made, appropriate post hoc tests were conducted as indicated in the figure legends. All statistical tests used were two-tailed. Experimenters were blind to experimental conditions during analysis.

---

**Figure S1.**
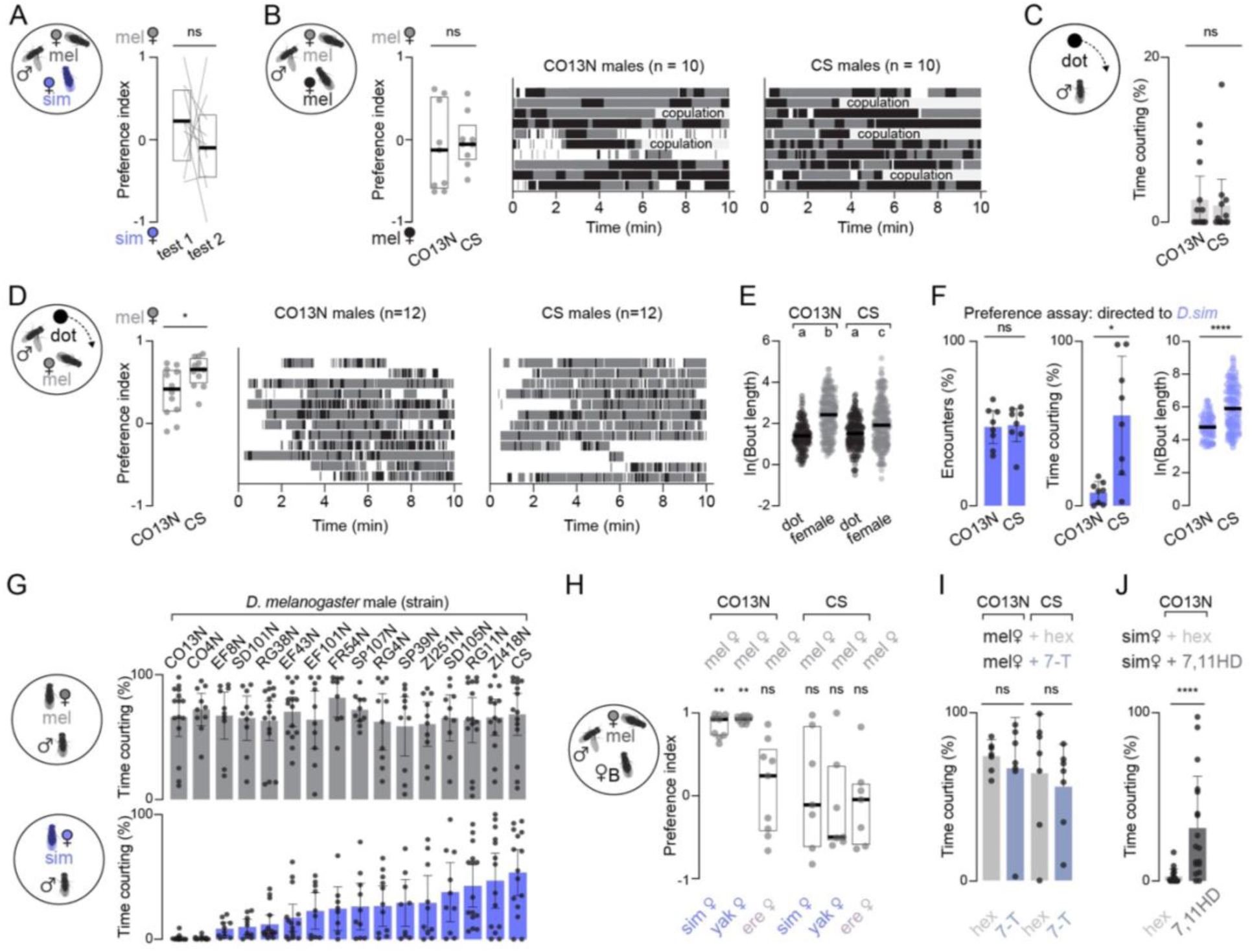
Male behavior implicates differential sensitivity to 7-T in establishing variation in strain preference. (**A**) Paired preference indices for CS males with a *D. melanogaster* and *D. simulans* female in two successive trials with individuals (lines), median (bold line), and inter-quartile range (IQR, box) shown. (**B**) Left: preference indices for CO13N and CS males paired with two equivalently reared *D. melanogaster* females with individuals (grey dots), median (bold line), and IQR (box) shown. Right: raster plots showing bouts of courtship over time toward each *D. melanogaster* female (grey or black) for CO13N and CS males, with copulations indicated. (**C**) Percent time spent courting a rotating dot projected onto the floor of the assay chamber (see Fig. 1D, Methods) for CO13N and CS males with individuals (grey dots), mean (bar), and 95% confidence interval (CI) shown. (**D**) Left: preference indices for CO13N and CS males paired with a *D. melanogaster* female (grey) and projected dot (black) with individuals (grey dots), median (bold line), and IQR (box) shown. Right: raster plots showing bouts of courtship over time toward the *D. melanogaster* female (grey) and the dot (black) for CO13N and CS males. (**E**) Natural log of bout lengths from (D) toward the dot and the female for CO13N and CS males with mean (bold line) shown. (**F**) Left: percent of all encounters (see Methods) with *D. simulans* female in the preference assay (Fig. 1C) for CO13N and CS males. Middle: percent time spent courting the *D. simulans* female for CO13N and CS males. Individuals (grey dots), mean (colored bars), and 95% CI (error bars) shown. Right: natural log of all bout lengths toward the *D. simulans* female in the preference assay (Fig. 1C) for CO13N and CS males with mean (bold line) shown. (**G**) Percent time spent courting a single *D. melanogaster* (grey, top) and *D. simulans* (blue, bottom) female for males of various strains with individuals (grey dots), means (colored bars), and 95% CI (error bars) shown. (**H**) Preference indices of CO13N and CS males paired with a *D. melanogaster* and a *D. simulans, D. yakuba,* or *D. erecta* female with individuals (grey dots), median (bold line), and IQR (box) shown. (**I**) Percent time that CO13N or CS males spent courting a single control *D. melanogaster* female treated with carrier solvent (hexane/hex, grey) or one perfumed with 7-T (blue-grey) with individuals (grey dots), mean (colored bars), and 95% CI (error bars) shown. (**J**) Percent time CO13N males spent courting a single control *D. simulans* treated with hexane (hex, light grey) or one perfumed with 7,11-HD (dark grey) with individuals (grey dots), mean (bars), and 95% CI (error bars) shown. ns, p > 0.05; *, p < 0.05; **, p < 0.01; ****, p < 0.0001, groups that are not significantly different from one another (p > 0.05) after multiple comparisons indicated by shared letter codes. Details of statistical analyses and sample sizes are given in Table S1.

**Figure S2.**
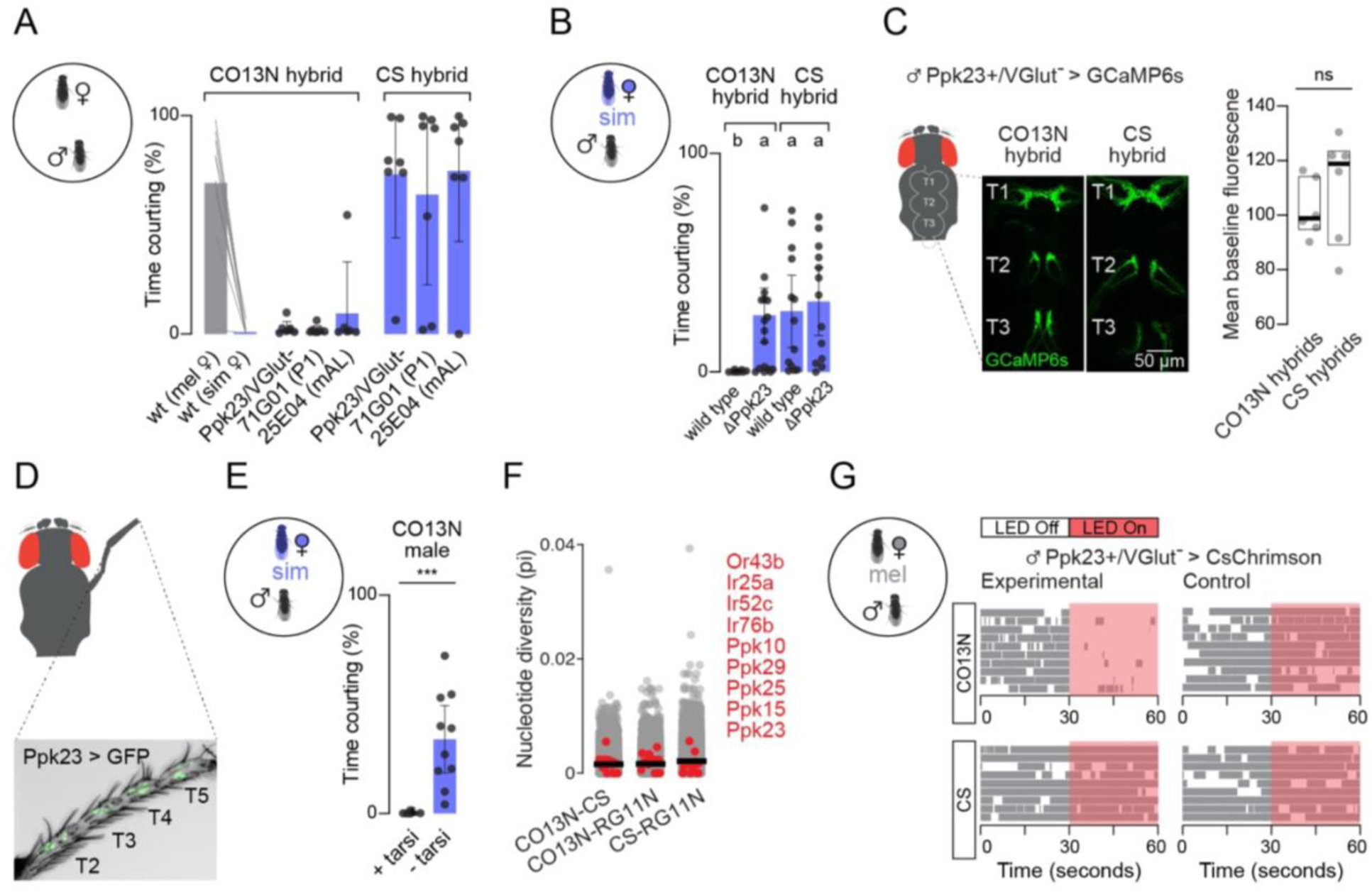
Cross-strain comparison of the sensory periphery. (**A**) Left: Paired courtship indices toward a *D. melanogaster* (grey) and *D. simulans* (blue) female for wild-type F1 hybrids generated by crossing CO13N and CS parents together (data where CO13N was the mother vs. where CS was the mother were not significantly different from one another and are combined in this figure). All other bars show courtship toward a *D. simulans* female for F1 crosses generated by crossing a wild-type selective CO13N or promiscuous CS female to the transgenic driver line indicated. (**B**) Percent time wild-type and ΔPpk23 null mutant CO13N and CS hybrid males spent courting a *D. simulans* female with means (bars), 95% confidence intervals (CI, error bars), and individuals (dots) shown. (**C**) Left: cartoon depicting the ventral nerve cord (VNC) of the male. Middle: 2-photon images of fluorescence from Ppk23-Gal4, VGlut-Gal80 > UAS-GCaMP6s in the VNC of selective CO13N (left) and promiscuous CS (right) hybrids. Right: mean baseline GCaMP6s fluorescence for drivers shown in (B). (**D**) Confocal image of a male foreleg with Ppk23-neurons marked by UAS-myr::GFP fluorescence. Tarsal segments are indicated. (**E**) Percent time that selective CO13N males courted a *D. simulans* female with intact forelegs or with foreleg tarsi (T2-T5) surgically removed. (**F**) Pairwise nucleotide diversity (pi) across all expressed transcripts (chemosensory gene transcripts highlighted in red and corresponding genes listed). Comparisons show sequenced RNA libraries generated from selective (CO13N) and promiscuous (CS, RG11N) or promiscuous (CS) and promiscuous (RG11N) foreleg samples. Mean pi indicated by black line. (**G**) Bouts of courtship toward a *D. melanogaster* female in the 30 seconds before and during optogenetic stimulus for Ppk23-Gal4, VGlut-Gal80 > UAS-CsChrimson CO13N and CS hybrid males shown in Fig. 2C raised on food with retinal (experimental) or without retinal (control).

**Figure S3.**
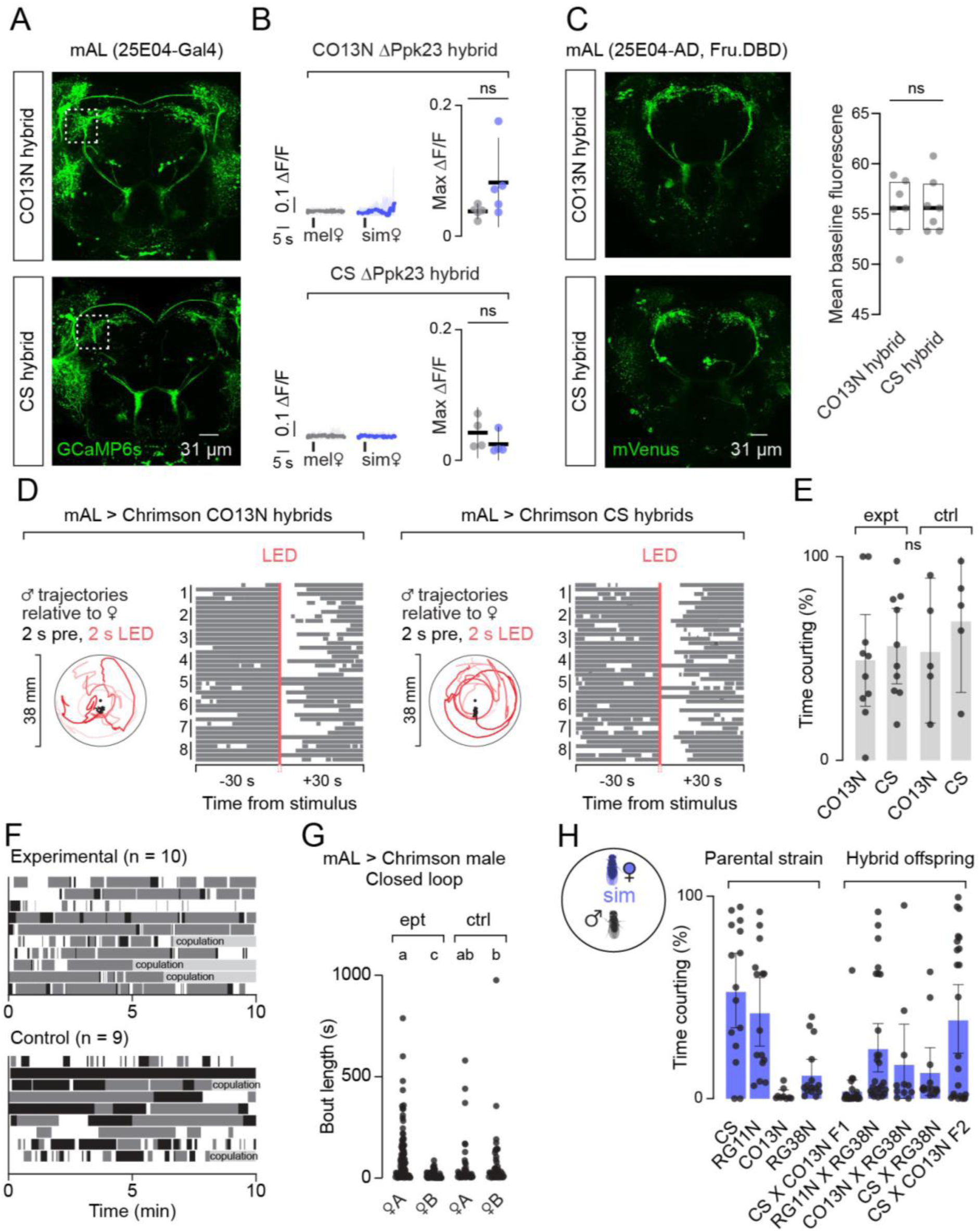
Cross-strain comparison of mAL neurons. (**A**) 2-P images of mAL neuron fluorescence in the brains of 25E04-Gal4 > UAS-GCaMP6s CO13N (top) and promiscuous CS (bottom) hybrids. White box indicates ROI for imaging mAL axons in the lateral protocerebral complex (LPC) shown in Fig. 4B. (**B**) mAL neuron responses in in ΔPpk23 mutant CO13N (top) and CS (bottom) hybrid males evoked by tapping the abdomen of a *D. melanogaster* (grey) or *D. simulans* (blue) female. Left panels: mean Δ*F*/*F_0_* traces (thick line) aligned to tap (black tick) with 95% CI shading. Right panels: maximum responses showing individual data points (dots), mean (line), and 95% CI (error bars). (**C**) Left: 2-photon images of fluorescence from 25E04-p64.AD, Fru-Gal4.DBD > UAS-GCaMP6s labeling mAL neurons in the brain of selective CO13N (top) and promiscuous CS (bottom) hybrids. Right: mean fluorescence from drivers shown in a standard ROI drawn over the LPC. (**D**) Left: Trajectories of representative mAL > UAS-CsChrimson CO13N (left) or CS (right) hybrid male in Fig. 4D plotted relative to the female’s position in the 2 seconds prior to (black) and 2 seconds during optogenetic stimulus (red) for each of the five stimuli. Right: Courtship bouts 30 seconds before, during, and 30 seconds after mAL optogenetic activation for all males in in Fig. 4D. (**E**) Percent time that experimental and control CO13N and CS hybrids spent courting over the full optogenetic assay in Fig. 4G; individuals (dots), mean (grey bars), and 95% CI (error bars) shown. (**F**) Courtship bouts toward female “A” (grey) and female “B” (black) in the closed-loop experiment (Fig. 4H–J) across the full 10 min assay; copulations marked. (**H**) Percent time promiscuous (CS, RG11N) and selective (CO13N, RG38N) parental strains and hybrid crosses spent courting a *D. simulans* female with individuals (dots), mean (blue bars), and 95% CI (error bars) shown.

**Table S1.** Statistical analyses by figure panel. Provided as a separate Excel file containing details of statistical analyses performed for each applicable figure and supplementary figure.

**Table S2.**
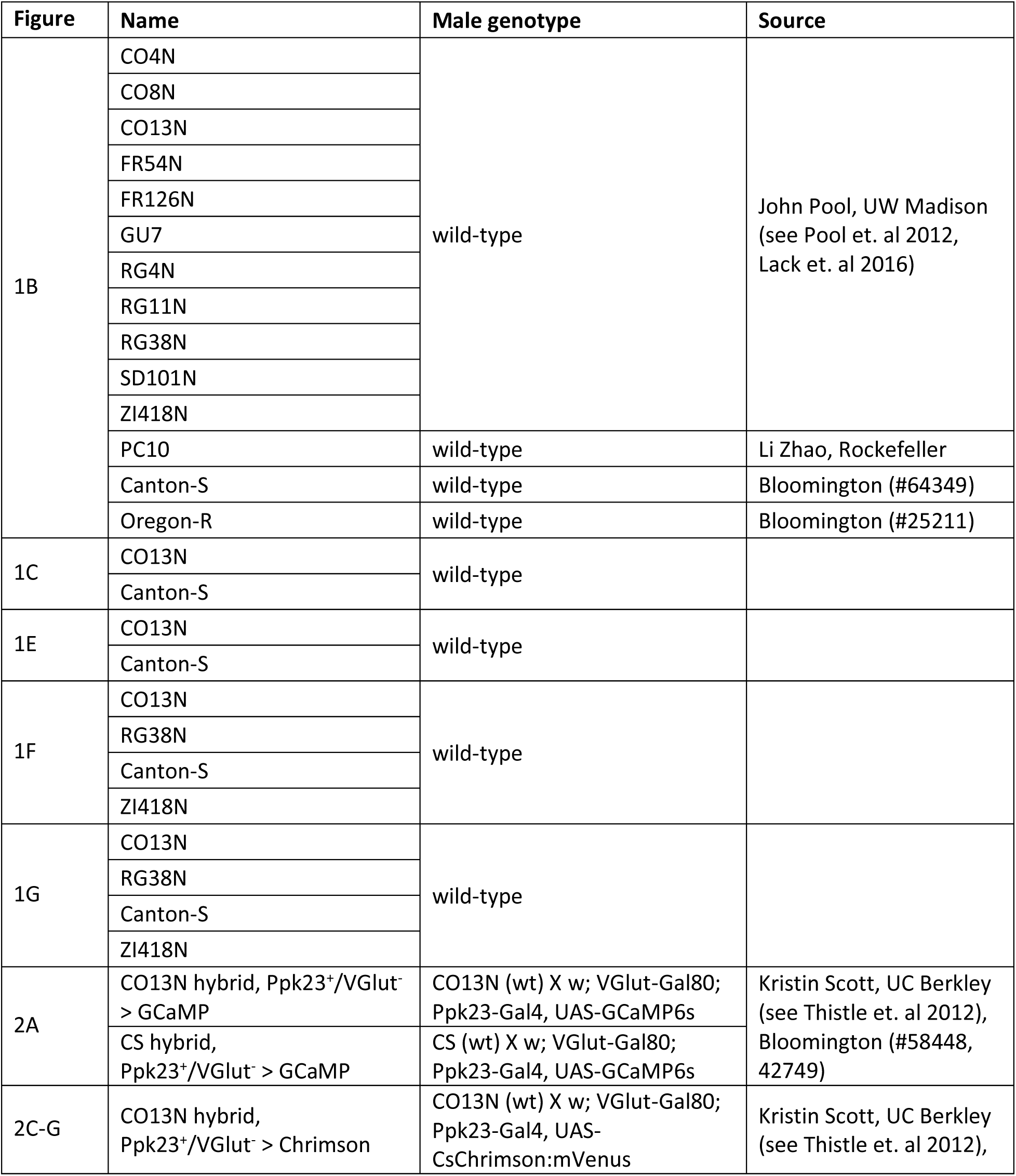
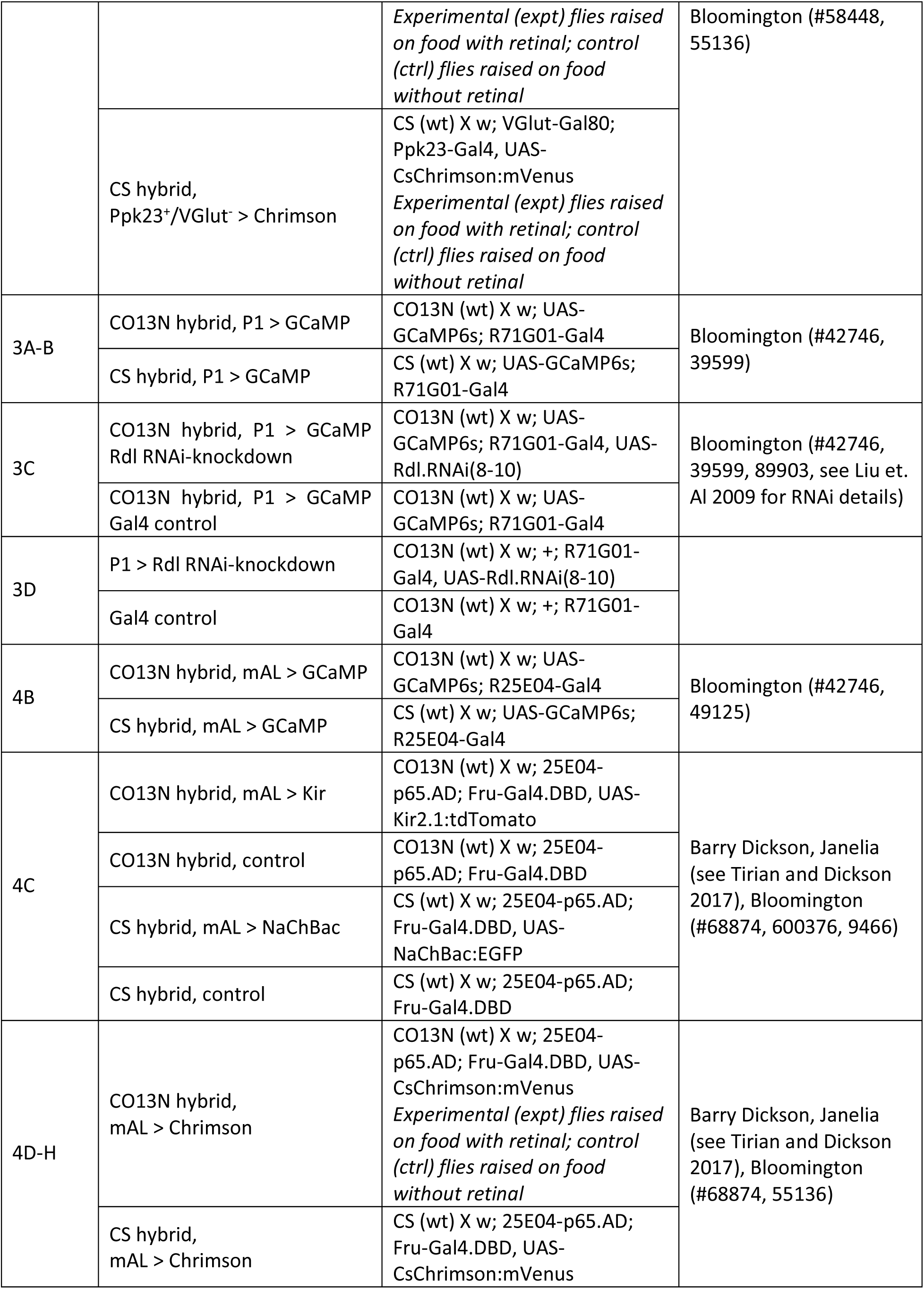
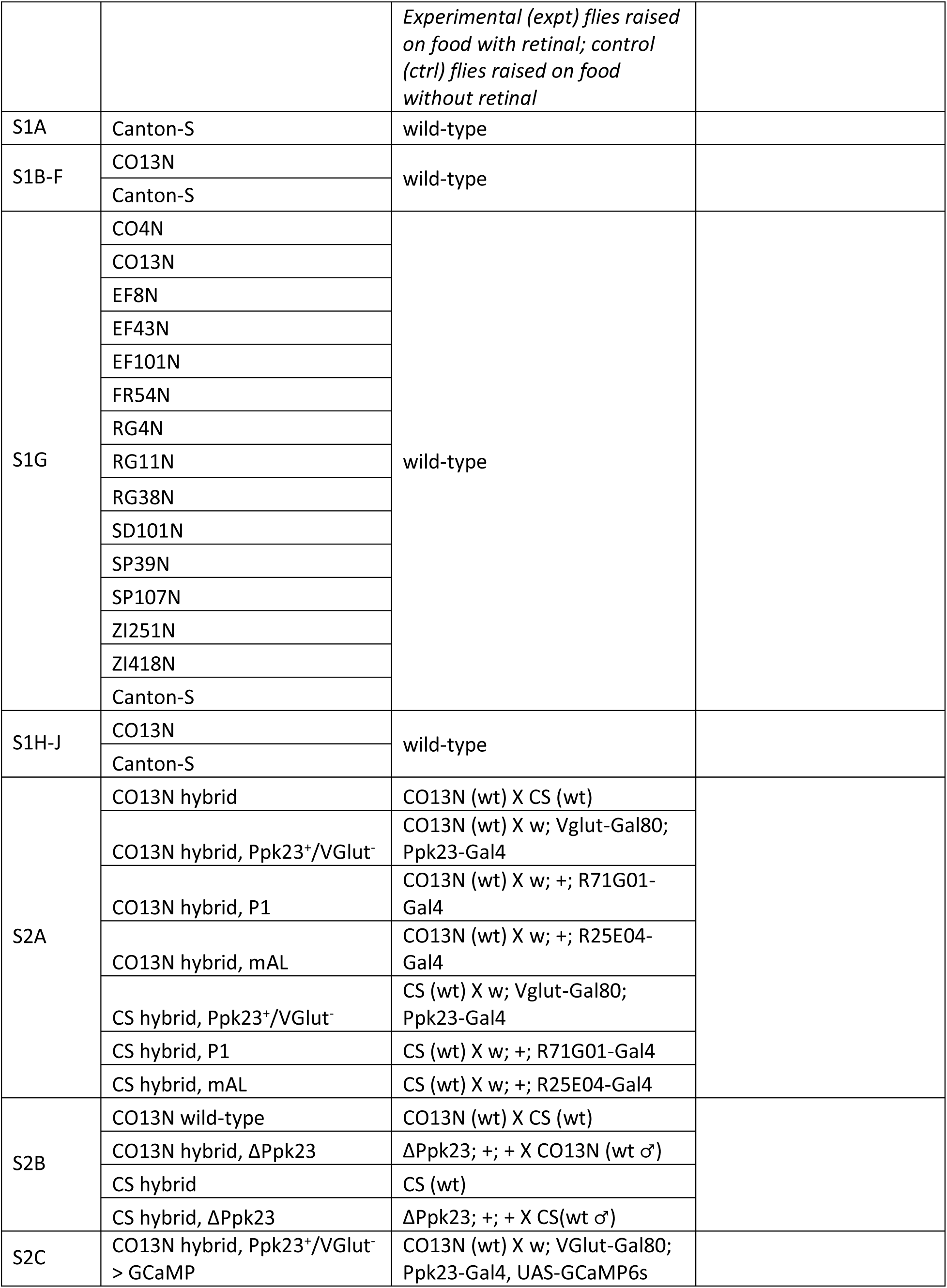
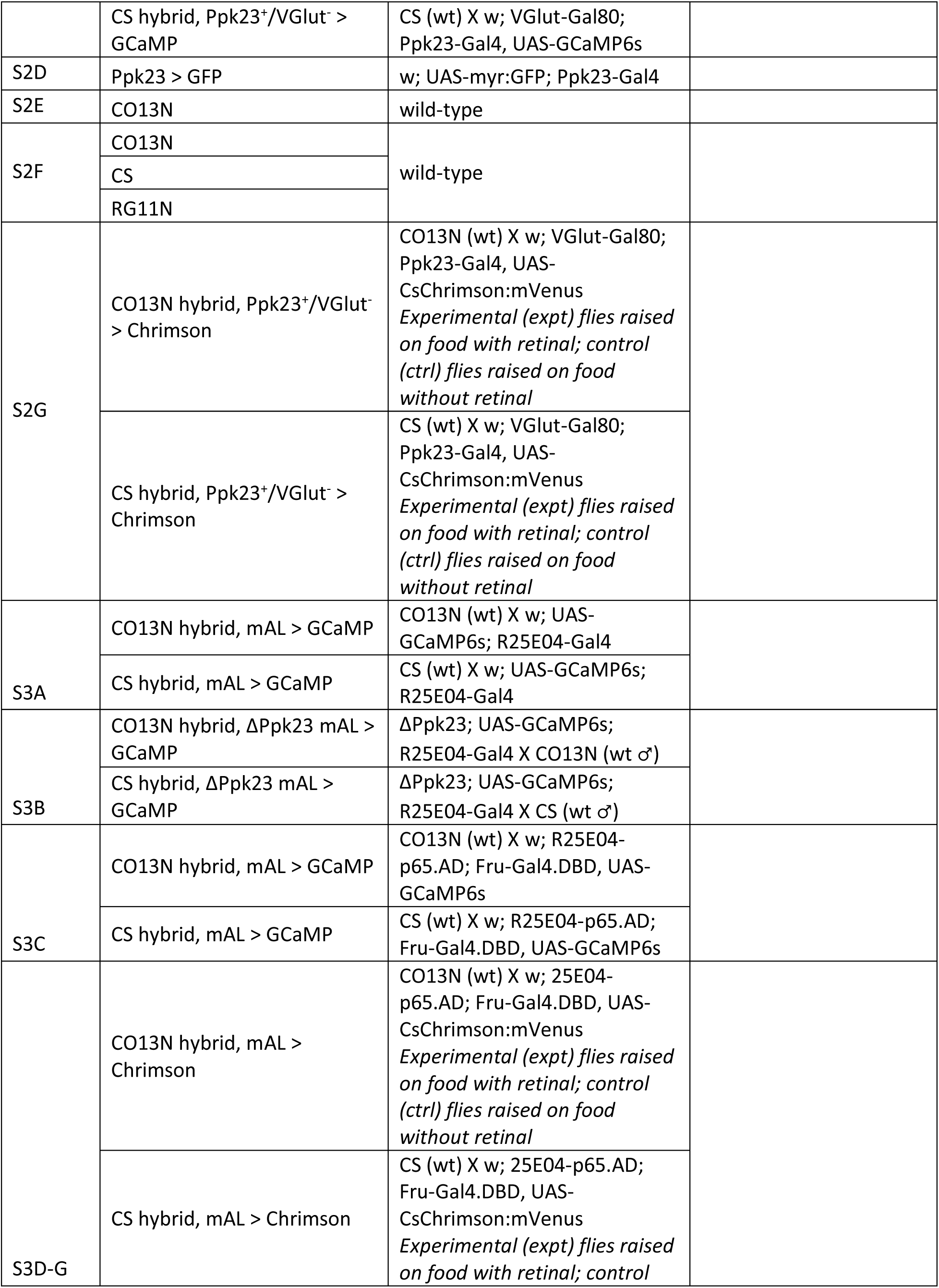
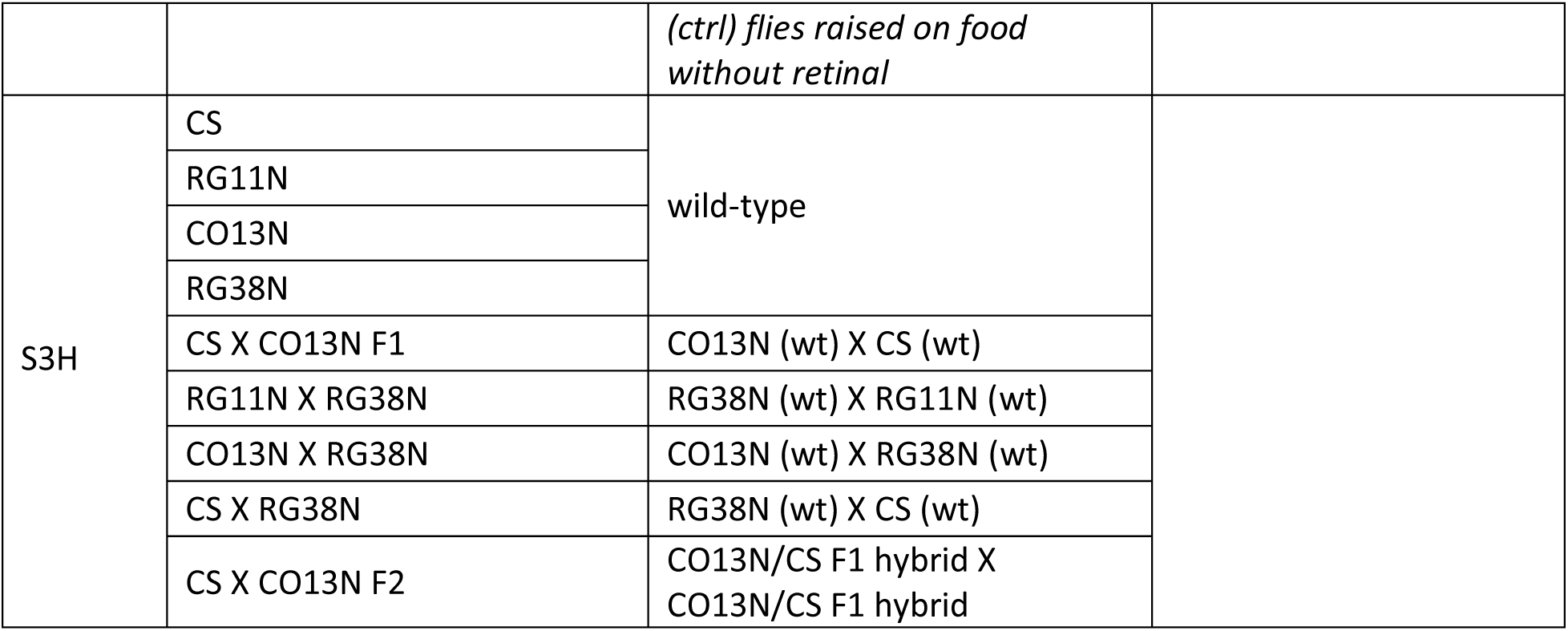
Fly genotypes and origin by figure panel. Associated genotypes of all males used in each figure section with their name as it appears in the text and their original source. *D. melanogaster* females were wild-type Canton-S. *D.simulans, D. erecta,* and *D. yakuba* females were wild-type.

**Movie S1. Behavior of a selective (CO13N) *D. melanogaster* male in preference assay, offered a conspecific and *D. simulans* female.** Representative video from a preference assay with identities of the participants indicated at the beginning of the clip and on subsequent frozen frames to aid with visual interpretation. Displayed at 2X speed.

**Movie S2. Behavior of a promiscuous (CS) *D. melanogaster* male offered a conspecific and D. simulans female.** Representative video from a preference assay with identities of the participants indicated at the beginning of the clip and on subsequent frozen frames to aid with visual interpretation. Videos are shown at 2X speed.

**Movie S3. Behavior of a selective (CO13N) male toward a projected dot alone and when paired with a conspecific female.** Sequential labeled video clips from assays in which a CO13N male interacts with a 3 mm dot projected onto the floor of the chamber alone and in the presence of a conspecific female. Videos are shown at 2X speed.

**Movie S4. Behavior of a promiscuous (CS) male toward a projected dot alone and when paired with a conspecific female.** Sequential labeled video clips from assays in which a CS male interacts with a 3 mm dot projected onto the floor of the chamber alone and in the presence of a conspecific female. Videos are shown at 2X speed.

**Movie S5. Optogenetic activation of 7T-sensing neurons in a selective hybrid male.** Representative video clips of CO13N hybrid males expressing CsChrimson in Ppk23⁺/VGlut⁻ sensory neurons, reared either on food with retinal (experimental) or without retinal (control). In all clips, males court a conspecific female and are then exposed to a 2 second red LED stimulus (LED-on indicated by a red circle in the upper left corner). Videos are shown at 2X speed.

**Movie S6. Optogenetic activation of 7T sensing neurons in a promiscuous hybrid male.** Representative video clips of CS hybrid males expressing CsChrimson in Ppk23⁺/VGlut⁻ sensory neurons, reared either on food with retinal (experimental) or without retinal (control). In all clips, males court a conspecific female and are then exposed to a 2 second red LED stimulus (LED-on indicated by a red circle in the upper left corner). Videos are shown at 2X speed.

**Movie S7. Optogenetic activation of mAL neurons in a selective hybrid male paired with two conspecific females.** Representative video clips of CO13N hybrid males expressing CsChrimson in mAL neurons (defined by the intersection of 25E04-p65.AD/Fru-Gal4.DBD), reared either on food with retinal (experimental) or without retinal (control). In all clips, males are paired with two conspecific females and exposed to a 2 second red LED stimulus (LED-on indicated by a red circle in the upper left corner). Videos are shown at 2X speed.

**Movie S8. Optogenetic activation of mAL neurons in a promiscuous hybrid male paired with two conspecific females.** Representative video clips of CS hybrid males expressing CsChrimson in mAL neurons (defined by the intersection of 25E04-p65.AD/Fru-Gal4.DBD), reared either on food with retinal (experimental) or without retinal (control). In all clips, males are paired with two conspecific females and exposed to a 2 second red LED stimulus (LED-on indicated by a red circle in the upper left corner). Videos are shown at 2X speed.

**Movie S9. Closed-loop ontogenetic activation of mAL neurons in a male paired with two conspecific females.** Representative video clips of CS hybrid males expressing CsChrimson in mAL neurons (defined by the intersection of 25E04-p65.AD/Fru-Gal4.DBD), reared either on food with retinal (experimental) or without retinal (control). In all clips, males are paired with two conspecific females and exposed to 500 ms second red LED stimulus (LED-on indicated by a red circle in the upper left corner) whenever real-time tracking algorithm registers him < 2mm and oriented within a 60-degree arc of the arbitrarily chosen fictive “*D. simulans*” female. Automated tracking of head/thorax/abdomen and the identity (male, blue; female, gold; fictive *D. simulans,* green) overlaid on video. Videos are shown at 2X speed.

## Notes

### Competing Interest Statement

The authors have declared no competing interest.

### Summary of Updates

Funding information on bioRxiv website updated.

